# Synaptotagmin 1 oligomerization via the juxtamembrane linker regulates spontaneous and evoked neurotransmitter release

**DOI:** 10.1101/2021.07.28.454225

**Authors:** Kevin C. Courtney, Yueqi Li, Jason D. Vevea, Zhenyong Wu, Zhao Zhang, Edwin R. Chapman

**Affiliations:** Howard Hughes Medical Institute and the Department of Neuroscience, University of Wisconsin, 1111 Highland Avenue, Madison, Wisconsin, 53705; Center for Bioanalytical Chemistry, Hefei National Laboratory for Physical Sciences at the Microscale, Department of Chemistry, University of Science and Technology of China, Hefei, P. R. China, 230027

## Abstract

Synaptotagmin-1 (syt1) is a Ca^2+^ sensor that regulates synaptic vesicle exocytosis. Cell-based experiments suggest that syt1 functions as a multimer, however biochemical and electron microscopy studies have yielded contradictory findings regarding putative self-association. Here, we performed dynamic light scattering on syt1 in solution, followed by electron microscopy, and used atomic force microscopy to study syt1 self-association on supported lipid bilayers under aqueous conditions. Ring-like multimers were clearly observed. Multimerization was enhanced by Ca^2+^ and required anionic phospholipids. Large ring-like structures (∼180 nm) were reduced to smaller rings (∼30 nm) upon neutralization of a cluster of juxtamembrane lysine residues; further substitution of residues in the second C2-domain completely abolished self-association. When expressed in neurons, syt1 mutants with graded reductions in self-association activity exhibited concomitant reductions in: a) clamping spontaneous release, and b) triggering and synchronizing evoked release. Thus, the juxtamembrane linker of syt1 plays a crucial role in exocytosis by mediating multimerization.

---

Synchronous release of neurotransmitters from presynaptic nerve terminals is triggered by the influx of Ca^2+^ ions via voltage gated Ca^2+^ channels (1–3). The rise in [Ca^2+^]i is sensed by the synaptic vesicle (SV) protein, synaptotagmin 1 (syt1) (4, 5), which interacts with anionic phospholipids (6–9) and SNARE proteins (10, 11) to trigger membrane fusion (12). Syt1 also plays additional roles in the regulation of the SV cycle, including: the inhibition of spontaneous release (clamping) (13–15), docking and priming (16, 17), and endocytosis (18–21). Although the importance of syt1 in the SV cycle is well established, whether and how syt1 performs each of these functions as a multimeric structure remains the subject of a long-standing debate.

Early work, using *Drosophila* as a model system, revealed intragenic complementation between several distinct mutant alleles of syt1 (13). This result strongly suggests that syt1 has separable functional domains and operates as a multimer *in vivo*. More recently, careful titration of mutant forms of syt1 (that were identified in human patients), in the presence of the wild-type protein, resulted in the potent dominant negative inhibition of SV exocytosis (22). This result further suggests that syt1 functions as an oligomer. However, the ability of syt1 to oligomerize, and the structure of these putative complexes, at both biochemical and ultrastructural levels, remains controversial.

SVs each contain approximately fifteen copies of syt1 (23), and multimerization of these monomers has been proposed to occur via two distinct mechanisms. The first proposed mode was constitutive and was mediated by determinants in the N-terminal region of the protein, which comprises residues 1-142 (Fig. 1A). Using density gradient centrifugation, native syt1 was reported to assemble into N-terminal-region mediated dimers (24) or tetramers in a non-denaturing detergent, CHAPS (4), but was also found to be largely monomeric in the absence of Ca^2+^ in Triton X-100 (25). N-terminal oligomerization was later refined in detergent-free conditions, and shown to occur within residues 1-96, which comprises a short intraluminal segment, a transmembrane domain (TMD), and the first sixteen residues of the juxtamembrane linker (residues 80-142) between the TMD and the first Ca^2+^-sensing motif, C2A (26).

**Fig. 1.**
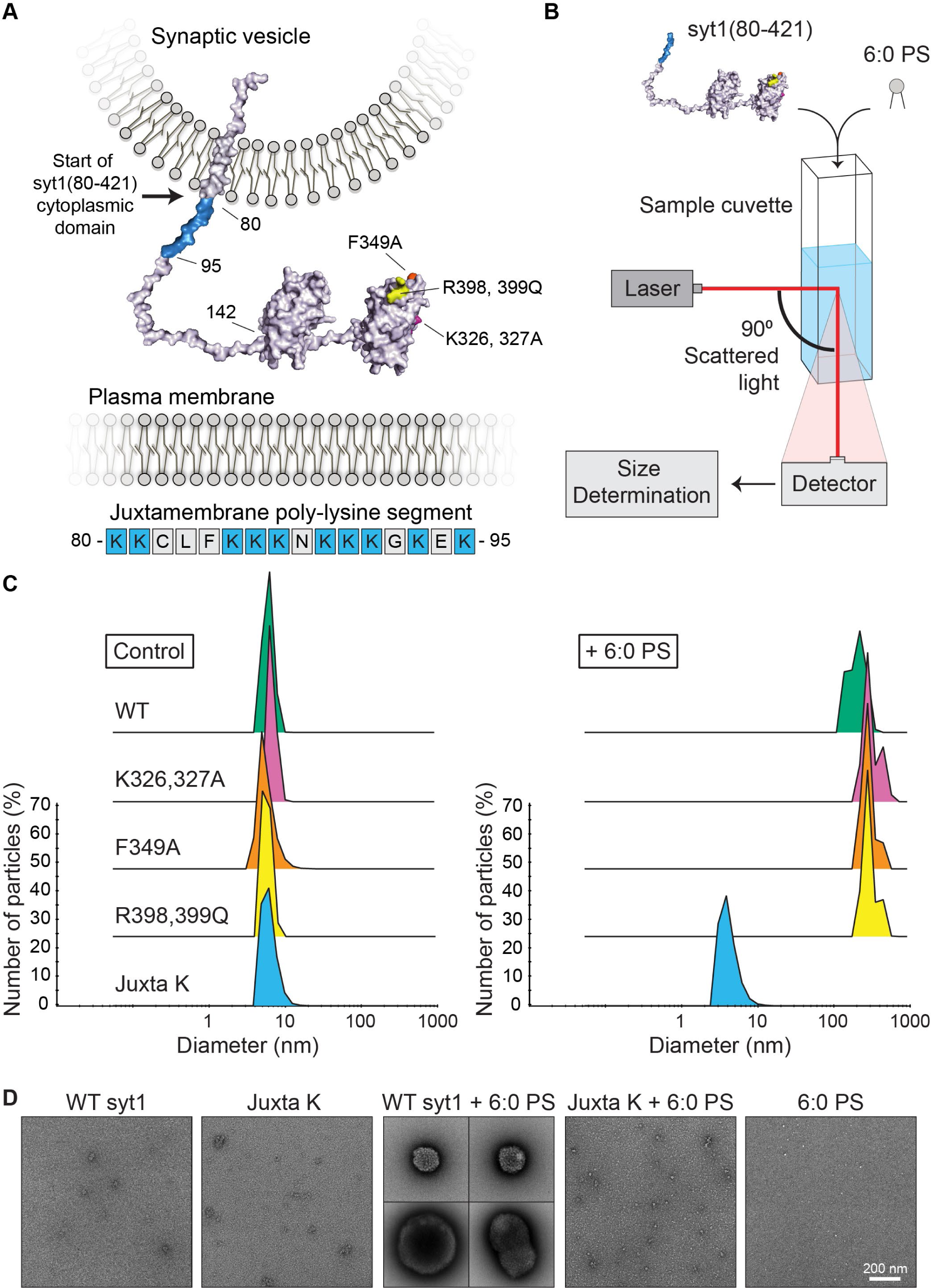
The complete cytoplasmic domain of syt1 forms large oligomers in solution upon addition of 6:0 PS. **A**) Structure of full-length syt1, embedded in a synaptic vesicle (SV) membrane, with relevant residue annotations. Structures of the C2-domains were derived from RCSB PDB 5t0r (C2A) and PDB 2yoa (C2B); other linker segments, derived from PBD 5w5c, were added. The syt1 oligomerization mutations tested in this study are color coded as follows: K326,327A, magenta; F349A, orange; and R398,399Q, yellow. The polylysine juxtamembrane region is emphasized in blue. The amino acid sequence (residues 80-95) of the syt1 juxtamembrane region immediately following the transmembrane domain. Lysine residues are shown in blue. **B**) Cartoon depiction of the DLS assay. **C**) Representative results of the DLS analysis showing the diameter of 2 µM WT and mutant syt1(80-421) with and without the addition of 100 µM 6:0 PS. Average diameters for each condition, performed in triplicate, are shown in Table 1. **D**) Representative EM images of WT and Juxta K mutant syt1(80-421) samples that were analyzed by DLS, with and without the addition of 6:0 PS. Scale bar represents 200 nm.

**Table 1.** Average diameter of syt1(80-421) in solution. DLS analysis of WT and mutant syt1(80-421) (2 µM) in aqueous buffer, with and without the addition of 100 µM 6:0 PS. Error represents standard deviation, derived from 3 independent experiments.

|  | WT | K326,327A | F349A | R398,399Q | Juxta K |
| --- | --- | --- | --- | --- | --- |
| Control | 6.0 ± 0.3 | 5.8 ± 1.3 | 5.1 ± 1.4 | 5.8 ± 0.7 | 5.4 ± 1.7 |
| + 6:0 PS | 215.7 ± 38.3 | 366.1 ± 22.2 | 321.3 ± 45.5 | 373.3 ± 7.9 | 5.1 ± 2.5 |

The second proposed mode of syt1 oligomerization is mediated by the C-terminal Ca^2+^- sensing domain, C2B (Fig. 1A). The syt1 C2B domain has been implicated in numerous modes of oligomerization that range from Ca^2+^ triggered homo-dimerization to the formation of large multimeric complexes (27–29). Reported oligomerization motifs within C2B include residues K326/327 (27), F349 (28) and R398/399 (29). However, even though self-association of purified, recombinant C2B has been abundantly described (25, 30), this protein fragment was found to tightly bind bacterial contaminants (31). These contaminants alter the biochemical properties of C2B, and thus complicate the interpretation of studies that did not include high salt washes and an RNAse/DNAse treatment to remove them during syt1 purification. Indeed, removal of these contaminants from C2B abolished its ability to self-associate in the presence or absence of Ca^2+^. However, in some studies, the addition of anionic phospholipid appeared to restore Ca^2+^-triggered oligomerization activity (29, 32–34).

Most recently, in a notable series of electron microscopy studies by Rothman, Krishnakumar, Volynski and colleagues, the cytoplasmic domain of syt1 formed ring-like oligomers on lipid monolayers when imaged via negative stain EM; these rings comprised >15 copies of syt1 (32). Interestingly, these rings were reported to be disrupted by Ca^2+^, and this dissolution was proposed to act as a mechanism to unclamp SV fusion (28, 32, 35–37). In contrast, other EM studies showed syt1 forms heptameric barrels in the presence of Ca^2+^ (34).

Due to the lack of consensus regarding the ability of syt1 to oligomerize, and the contradictory reports regarding the structure of these putative oligomers, we re-examined this question while adhering to physiological conditions that required minimal manipulation of the samples. To this end, we performed dynamic light scattering (DLS) on syt1 in solution, followed by electron microscopy (EM), and atomic force microscopy (AFM) imaging of syt1 on supported lipid bilayers (SLB). Together, these *in vitro* experiments served as a platform to screen for putative oligomerization motifs that were then functionally tested in neurons. Importantly, these *in vitro* experiments were conducted under aqueous conditions. Moreover, we used the complete cytoplasmic domain of the protein (residues 80-421) (Fig. 1A). This contrasts with most of the earlier *in vitro* work which utilized truncated forms of the cytoplasmic domain of syt1 (i.e. residues 96-421 or 143-421) that lacked either the highly cationic portion of the juxtamembrane region or the entire juxtamembrane segment (Fig. 1A). By DLS and EM, we observed that anionic lipids cause syt1(80–421) to assemble into large clusters. Moreover, AFM imaging found that syt1(80–421) forms clusters, large ring-like structures, and large patches on phospholipid bilayers in an anionic lipid dependent manner; formation of these multimeric structures on phospholipid bilayers was enhanced by Ca^2+^. Mutagenesis revealed that lysine residues in the juxtamembrane linker are crucially important for syt1 self-association. AFM imaging revealed that additional mutations, in the C2B domain, further diminished the ability of syt1 to self-associate. In neurons, the ability of syt1 to multimerize was well correlated with its ability to clamp spontaneous release and to trigger and synchronize evoked release. Hence, the juxtamembrane linker enables syt1 to assemble into a multimer, and this multimerization plays a key role in the regulation of SV exocytosis.

## RESULTS

### The syt1 cytoplasmic domain forms multimeric structures, in solution, upon binding anionic phospholipids

The structure of syt1 is depicted in Fig. 1A. The N-terminal region comprises a short luminal domain, a single TMD and a juxtamembrane linker that is followed by the tandem C2- domains (24). Within this juxtamembrane linker lies a sequence containing ten positively charged residues directly after the TMD (Fig. 1A). A key feature of the experiments reported here is that we used the intact cytoplasmic domain of syt1, residues 80-421, which includes this cationic juxtamembrane segment. Again, this contrasts with the majority of published work describing syt1 biochemistry and oligomerization because those reports were based on shorter fragments (residues 96-421 or 143-421, both lacking the polybasic region) (28–30, 32).

We first assessed the self-association properties of syt1 by performing DLS in aqueous media (Fig. 1B). Each of the two C2 domains of syt1 are approximately 2.5 x 5 nm (*SI Appendix*, Fig. S1A). We reasoned that if syt1 assembled into a multimeric structure, the average hydrodynamic diameter of the pure protein should exceed these monomeric dimensions. When suspended in physiological, aqueous media, pure syt1(80–421) was found to have an average diameter of approximately 5 nm, suggesting it is indeed monomeric (Fig. 1C). Syt1 is known to function by binding anionic lipids, namely phosphatidylserine (PS) and PIP2 (6–9). We therefore examined how anionic lipids would influence multimerization. Remarkably, we found that the addition of a soluble, short chain anionic lipid, 6:0 PS, caused syt1 to assemble into ∼ 250 nm structures (Fig. 1C). In contrast, syt1 remained monomeric in the presence of a non-acylated variant (phospho-serine) (*SI Appendix*, Fig. S1B) and 6:0 PS alone failed to generate detectable DLS signal (data not shown). This demonstrates that syt1 self-association is triggered upon binding anionic phospholipids, mediated by electrostatic and hydrophobic interactions. We then proceeded to image the DLS samples by negative-stain EM. In line with the DLS, EM imaging found large clusters in the WT syt1(80–421) + 6:0 PS sample, whereas protein alone and 6:0 PS alone samples were devoid of these large structures (Fig. 1D).

Since SVs have, on average, 15 copies of syt1 (23), we do not expect these large structures to exist *in vivo*. However, these findings show that the cytoplasmic domain of syt1 has the ability to self-associate in the presence of anionic lipids. As such, we reasoned that analyzing the size of these supraphysiological multimeric structures, in conjunction with site-directed mutagenesis, would enable us to map the determinants that mediate lipid-dependent homomeric interactions under aqueous conditions.

### Syt1 self-association persists after substitution of residues in the C2B domain

To gain insight into the structural elements of syt1(80–421) that mediate self-association, we mutated a number of residues that have previously been reported to regulate syt1 oligomerization through the C2B domain and performed DLS analysis. These mutant forms of syt1 encompass: K326,327A (27, 34), a positively charged region that is also responsible for Ca^2+^- independent PIP2 binding activity (7); F349A, which was reported to disrupt the ring structures observed by negative stain EM (28, 36); and R398,399Q, implicated in binding SNAREs and C2B self-association via back-to-back dimerization (29). All three sets of mutations failed to disrupt multimerization under our experimental conditions (Fig. 1C and Table 1).

### Lysine residues in the juxtamembrane linker of syt1 regulate self-association

As outlined in the Introduction, syt1 was first thought to self-associate via determinants in the N-terminus of the protein (4, 26). Within this putative oligomerization region, the juxtamembrane linker was shown to mediate Ca^2+^-independent interactions with membranes (38). The juxtamembrane linker contains a segment (between residues 80-95) in which ten of the first sixteen residues after the transmembrane domain are lysines (Fig. 1A). This cationic region is poorly characterized as residues 80-95 are commonly excluded from recombinant preparations of the syt1 soluble domain, perhaps due to increased difficultly in purification (see Methods). We hypothesized that these charged residues mediate interactions with anionic phospholipids to promote oligomerization (4, 26, 38). To assess this possibility, we neutralized the juxtamembrane lysines (Juxta K) (*SI Appendix*, Fig. S2) and, again, performed DLS on syt1(80–421) with and without 6:0 PS. In contrast to WT protein, we found that 6:0 PS failed to trigger self-association of the Juxta K variant; Juxta K syt1 remained monomeric in solution (Fig. 1C and Table 1). This DLS result was further validated by EM imaging, which found that the Juxta K + 6:0 PS sample completely lacked large protein-lipid clusters (Fig. 1D).

### The syt1 cytoplasmic domain forms multimeric structures on supported lipid bilayers

Although DLS and EM analysis served as an efficient screen that enabled us to uncover Juxta K mediated syt1 self-association, this system is accompanied by non-physiological caveats. To validate the DLS and EM results, we developed a second *in vitro* assay to examine if the syt1 cytoplasmic domain self-associates on the surface of phospholipid bilayers. For this, we generated SLBs and performed AFM imaging while aiming to mimic native, physiological conditions (Fig. 2A). This AFM strategy combines the distinct advantages of both DLS and EM by enabling high sensitivity experiments, with single particle resolution, in aqueous media. As preliminary test of our AFM approach, we incubated 1 µM syt1(80–421) with a SLB (DOPC/DOPS/PIP2, 72:25:3) for 6 hours in the absence of Ca^2+^. Under these aqueous conditions, we observed the formation of large numbers of ring-like structures formed by syt1 on the lipid bilayer surface (*SI Appendix*, Fig. SA). A zoomed-in three-dimensional view of a representative structure is also shown (*SI Appendix*, Fig. S3B). Overall, the diameter of these structures (i.e. the distance between the two highest points on both sides in a cross-section) was commonly greater than 100 nm, suggesting that they are formed by a high copy number of syt1(80–421).

**Fig. 2.**
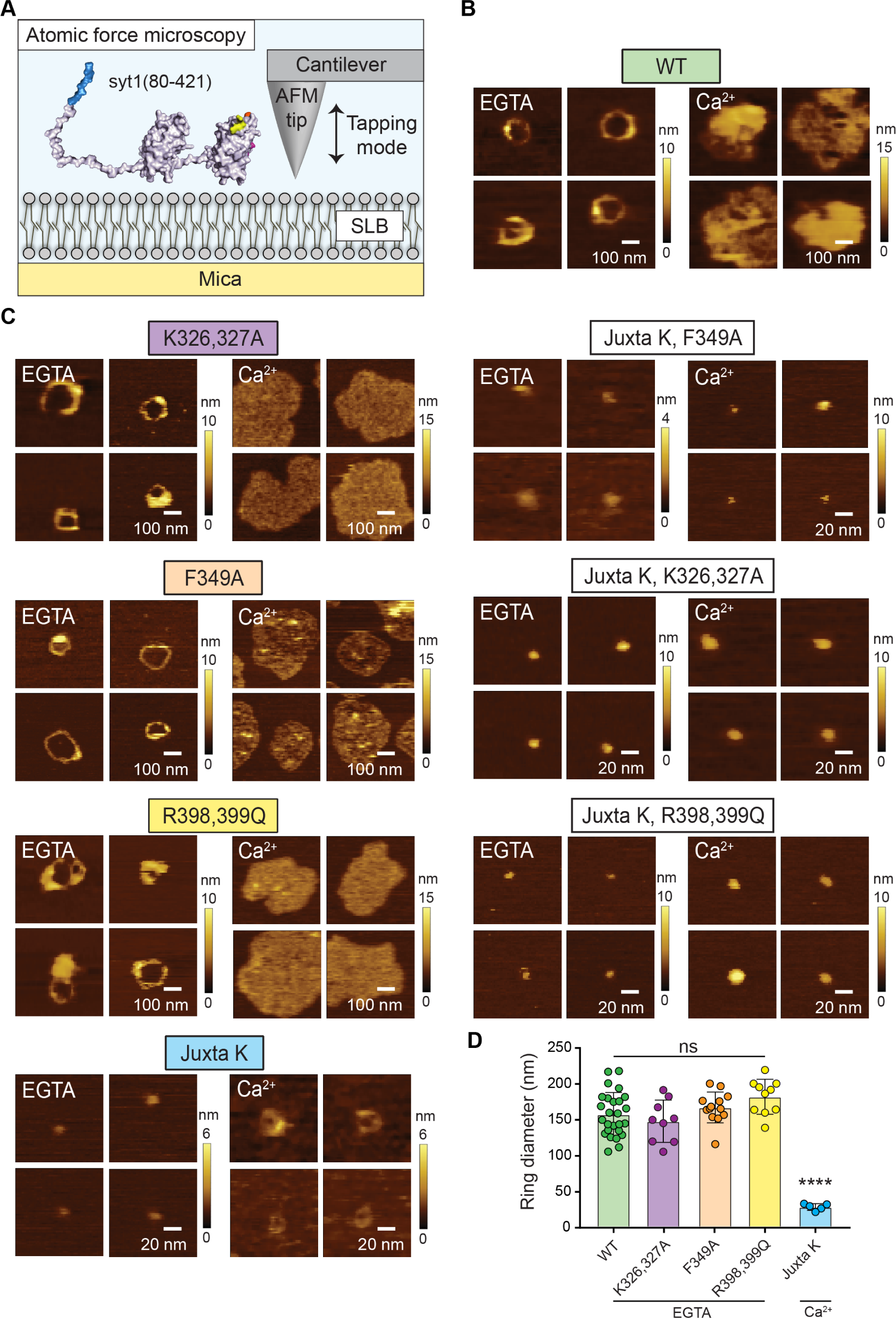
AFM imaging of WT and mutant syt1(80-421) on supported lipid bilayers. **A**) Cartoon depiction of the AFM experimental conditions. **B**) Representative AFM topographical images of 1 µM WT syt1(80-421) on supported lipid bilayers (72% DOPC, 25% DOPS, 3% PIP2) in the presence (1 mM) and absence (0.5 mM EGTA) of Ca^2+^. **C**) Representative AFM topographical images of the syt1(80-421) mutants under the same conditions as the WT samples. The protein concentration was 1 µM in all cases except for the R398,399Q sample, where a concentration of 300 nM was used to better visualize ring formation. Pixel size: left column 9.8 nm/px, right column 2.0 nm/px. **D**) Ring diameters for WT syt1(80-421) (green), K326,327A (purple), F349A (orange), and R398,399Q (yellow) in the absence of Ca^2+^, and of Juxta K (blue) in the presence of Ca^2+^. For the Juxta K sample in Ca^2+^, 41.6% of the observed structures were rings, while the remainder were particles. Additional C2B mutations, in the Juxta K background, abolished ring formation so these samples were excluded from ring analysis. **** Denotes p-values < 0.0001 for Juxta K, compared to all other conditions. For WT, F349A, K326,327A, R398,399Q and Juxta K, the number of rings measured were: 26, 13, 9, 10 and 5 for each condition from 3, 2, 2, 2, and 2 sets of analyses using independent SLBs, respectively. Error bars represent SEM.

Importantly, the syt1 rings that we observed by AFM are filled with lipid in the center, as determined by lateral height profiles in conjunction with an analysis of surface roughness inside and outside the rings (*SI Appendix*, Fig. S3B and S3C). These findings indicate that these structures assemble on the surface of intact bilayers. We also occasionally observed structures that formed around defects (holes) in the SLB (*SI Appendix*, Fig. S4A). These structures resemble protein-decorated holes that form on SLBs after treatment with pore-forming proteins, such as Bax (Salvador-Gallego et al., 2016). However, since syt1(80–421) does not form large pores in bilayers (*SI Appendix*, Fig. S4B), we believe the effects of pore-forming proteins, like Bax, on SLBs are distinct from the ring-like structures that we observed. Indeed, upon further examination, we found that defects in the SLB are stable over time (*SI Appendix*, Fig. S4C), while the number of ring-like structures increase dramatically with time (*SI Appendix*, Fig. S4D). Hence, syt1 ring formation does not require membrane defects, and these ring-like structures represent bona fide syt1 multimers on the SLB surface. Notably, since the properties of the structures that associate around membrane defects are dominated by the size and shape of the defect itself, rather than the multimerization properties of syt1, we excluded structures with an interior hole or defect in the bilayer from all analyses.

### Syt1 self-association requires anionic phospholipids and is enhanced by Ca^2+^

After establishing that circular multimerization (i.e. ring formation) on the bilayer surface was robust and reproducible by AFM imaging, we set the incubation time of syt1(80–421) with the SLB to 20 minutes in all subsequent trials. In the absence of Ca^2+^ (0.5 mM EGTA), at increasing protein concentration, protein structures on the SLB transitioned from particles (50 nM) to rings (1 µM), and from rings to patches (3 µM) (*SI Appendix*, Fig. S5 and S6; Table S1). The sensitivity of the multimeric structures to protein concentration in our AFM analysis, and the variation in the morphology of these multimers, suggest structural plasticity in the multimerization process. In comparison, in 1 mM free Ca^2+^, rings and patches still formed, but both classes of multimers formed at lower protein concentrations as compared to the Ca^2+^-free condition (*SI Appendix*, Fig. S6; Table S1). The effect of Ca^2+^ was also consistent across a range of Ca^2+^ concentrations (*SI Appendix*, Fig. S7A). The above findings show that all forms of syt1 self-assembly: particles, rings and patches, can occur in the absence of Ca^2+^, and that addition of Ca^2+^ facilitates multimerization. To further confirm this Ca^2+^-dependent enhancement, we tested a syt1 Ca^2+^ binding mutant, syt1^4N^(80–421), in which two acidic Ca^2+^ ligands in each C2-domain were mutated to neutral residues, thus abolishing Ca^2+^ binding activity (39, 40). Ring-like structures were still observed, but Ca^2+^ failed to enhance further assembly (*SI Appendix*, Fig. S7B). The precise mechanism by which Ca^2+^ promotes syt1 self-association is not yet known, but likely involves conformational changes that alter the relative orientation of its tandem C2-domains within the multimer (15). In addition, Ca^2+^ might also facilitate self-assembly by increasing the local syt1 concentration on the bilayer.

To examine the influence of the SLB phospholipid composition on syt1 self-association, we omitted PS and PIP2 in our protein-lipid interaction tests of syt1(80–421). We found that anionic phospholipids were required for syt1 multimers to assemble on the bilayer (*SI Appendix*, Fig. S8A). Next, to validate that phospholipid binding promotes syt1 self-association, we deposited syt1(80–421) onto a bare mica surface (lipid free), and again studied its morphology under aqueous conditions (*SI Appendix*, Fig. S8B). When no lipid was present, syt1(80–421) molecules appear as dispersed particles with similar dimensions in both EGTA and Ca^2+^ conditions.

### Lysine residues in the juxtamembrane linker of syt1 are essential for large ring formation on supported lipid bilayers

As described above, our DLS and EM analysis revealed that syt1 multimerization, in response to lipid binding, is governed by the Juxta K region. We revisited all the syt1 mutants characterized by DLS to assess their respective impacts on multimerization on phospholipid bilayers. The WT and mutant syt1(80–421) structures that were analyzed by AFM and the associated lateral height profiles are shown in *SI Appendix*, Fig. S9-S13. Our AFM analysis found the K326,327A, F349A and R398,399Q mutations have no effect on the formation of large multimeric structures, similar to the DLS result (Fig. 2C). However, also in agreement with our DLS and EM results, we found that neutralizing the juxtamembrane lysine residues (Juxta K) dramatically disrupted the formation of large rings on the SLBs (Fig. 2C). Interestingly, although no large multimeric structures were present with 1 µM Juxta K in 0.5 mM EGTA, smaller ∼30 nm ring structures were formed in the presence of 1 mM free Ca^2+^ (Fig. 2C and *SI Appendix*, Fig. S13). The distinct diameters of the small and large rings (Fig. 2D) suggest that the two populations of syt1(80–421) multimers form by different mechanisms. Notably, the smaller rings have dimensions that are comparable to the rings reported by Rothman, Krishnakumar, Volynski and colleagues (28, 32, 35, 36). In stark contrast, however, the small rings observed by AFM in aqueous buffer only appeared in the presence of Ca^2+^, whereas the rings observed by negative stain EM were dispersed by Ca^2+^ (32).

### Concurrent mutations in two distinct regions of syt1 abolish self-association

In an effort to further disrupt syt1 self-association (small rings), we analyzed all of the above C2B mutations in a Juxta K mutant background. Remarkably, only featureless particles were observed in the AFM images of Juxta K + K326,327A, Juxta K + F349A, and Juxta K + R398,399Q (Fig. 2C). These findings reveal that syt1 multimerization is regulated by a complex interplay between the juxtamembrane region and the C2B-domain of syt1, with the juxtamembrane region serving as the primary determinant. These data suggest that the C2B mutations may have had subtle effects on self-association that escaped detection when analyzed in an otherwise WT background (Fig. 2C).

### Syt1 self-association mutants are targeted to synapses with the correct topology

Having established that the complete cytoplasmic domain of syt1 forms homo-multimers under aqueous conditions on lipid bilayers, we sought to determine the functional relevance of this interaction by conducting cell-based experiments. We first determined whether each of the mutants described above were properly targeted to SVs in cultured syt1 KO mouse hippocampal neurons. Floxed syt1 was disrupted using Cre recombinase, followed by re-expression of WT or each mutant form of the protein using lentiviral transduction; expression was monitored via immunoblot (Fig. 3A and *SI Appendix*, Fig. S14; we note that juxtamembrane lysine mutations (Juxta K) reduced the mobility of both recombinant and neuronally expressed syt1 on SDS-PAGE gels). We observed that WT and each mutant form of syt1 were highly co-localized with the SV marker synaptophysin (Fig. 3B and 3C). Notably, the syt1 Juxta K mutant lentiviral expression construct preserved the WT K80 and K81 residues directly following the TMD to ensure proper syt1 topology (*SI Appendix*, Fig. S2) (41). To further confirm targeting to SVs, we conducted pHluorin experiments and found that upon stimulation and exocytosis, all constructs rescued the reduction in the time-to-peak that is characteristic of syt1 KO neurons (19) (Fig. 3D and 3E). Moreover, all but the F349A mutant rescued the kinetic defect in endocytosis that occurs in the KO (19) (Fig. 3F). The unexpected inability of F349A to rescue SV recycling will be addressed in a future study. In summary, these findings demonstrate that all constructs are targeted to SVs with the same topology as the WT protein. However, the low time resolution of the pHluorin measurements sharply limits what can be learned about excitation-secretion coupling, so we next turned to high-speed glutamate imaging and electrophysiology experiments to address the impact of the self-association mutants on evoked and spontaneous neurotransmitter release.

**Fig. 3.**
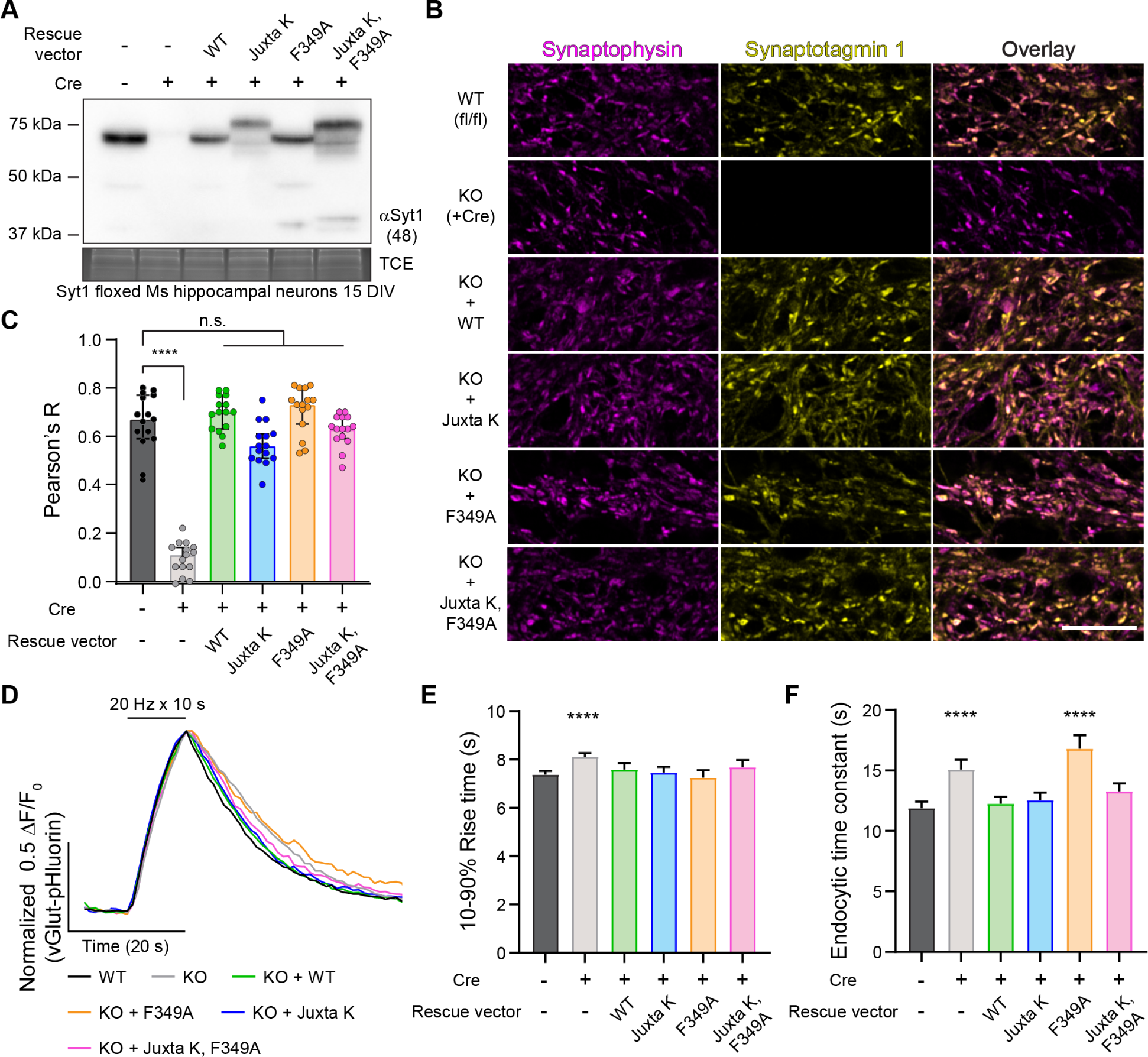
WT and mutant syt1 constructs are efficiently targeted to SVs. **A**) Representative anti-syt1 immunoblots of WT, syt1 KO (generated using a Cre virus), and KO neurons expressing WT syt1, Juxta K mutant syt1, F349A mutant syt1, and the double Juxta K + F349A mutant syt1, in mouse hippocampal neurons at 15 DIV. **B**) Representative super-resolution fluorescent ICC images from mouse hippocampal neurons at 21 DIV. Images of WT, syt1 KO (+Cre), and KO neurons expressing WT syt1, Juxta K mutant syt1, F349A mutant syt1, and the double Juxta K + F349A mutant syt1, stained with anti-synaptophysin (magenta) and anti-syt1 (yellow) antibodies; in the last column these signals are merged. **C**) Bar graph of the Pearson’s correlation coefficient (PCC) statistic R. Values were obtained using ROIs from whole fields of view and JaCoP for ImageJ (60). Plot of the median Pearson’s R +/- 95% Cl; n = 15 for each condition from three trials. These data were not normally distributed; **** denotes p-value < 0.0001 between WT and syt1 KO, no other conditions were statistically significantly different from WT using Kruskal-Wallis test with Dunn’s correction. Full statistics are included in Fig. 3-source data. **D**) Partially cropped, averaged, and normalized vGlut1-pHluorin traces from each indicated condition obtained using wide field fluorescence, imaged once a second for 120 seconds. Twenty Hz field stimulation began at t = 10 s (200 action potentials) in 14 DIV hippocampal mouse neurons. **E**) vGlut1-pHluorin 10%-90% peak rise times were plotted. Values are median with 95% CI; 500 to 700 ROIs (n) were analyzed from three separate trials. These data were not normally distributed’ **** denotes p-value < 0.0001 between WT and syt1 KO, no other conditions were statistically significantly different from WT using Kruskal-Wallis test with Dunn’s correction. **F**) vGlut1-pHluorin decay times represented as time constants determined from fitting data to a single exponential function. Values are median with 95% CI; 500 to 700 ROIs (n) measured from three separate trials. These data were not normally distributed; **** denotes p-value < 0.0001 between WT and syt1 KO, and WT and F349A mutant rescue, no other conditions were statistically significantly different from WT using Kruskal- Wallis test with Dunn’s correction.

### Syt1 self-association is essential for driving and synchronizing evoked SV exocytosis

In the next series of experiments, we used iGluSnFR (42) to monitor the evoked release of glutamate from SVs, triggered by single action potentials, in WT, syt1 KO, and syt1 KO neurons rescued with each of the constructs detailed above. From the raw traces (Fig. 4A), and from histograms that were created by binning the peak iGluSnFR signal (ΔF/F0) versus time (Fig. 4B), it was evident that loss of syt1 abolished rapid synchronous release; only slow asynchronous release was detected. Fast release was completely rescued by WT syt1 and the F349A mutant. In contrast, the Juxta K mutant, alone and combined with the F349A mutant, only partially rescued fast release (Fig. 4A and 4B). Examination of the average traces showed that expression of the Juxta K mutant resulted in a 63 ± 1.7 % reduction in the peak amplitude of the iGluSnFR signal, and this was exacerbated by adding the F349A mutation, resulting a 79 ± 1.7 % reduction in the signal (Fig. 4C, 4D and *SI Appendix*, Table S2). To visualize the influence that these mutations have on the balance of synchronous vs asynchronous release, we plotted the normalized cumulative frequency distributions for WT, and each mutant, as a function of time. From this analysis, the ability of syt1 to synchronize release is readily apparent (Fig. 5A); the synchronous fraction of total release observed over the image series was then extracted and is plotted in Fig. 5B. Substitution of F349 slightly reduced synchronization, but this did not reach significance in our experiments (but see ref. (37)). However, the Juxta K mutations, which strongly affected homo-multimerization, resulted in a marked reduction in the ability of syt1 to synchronize release. Moreover, this effect was even greater when the Juxta K mutant also included the F349A mutation (Fig. 5). Hence, there is a correlation between the ability of mutations to impair syt1 self-association and to reduce and desynchronize evoked release.

**Fig. 4.**
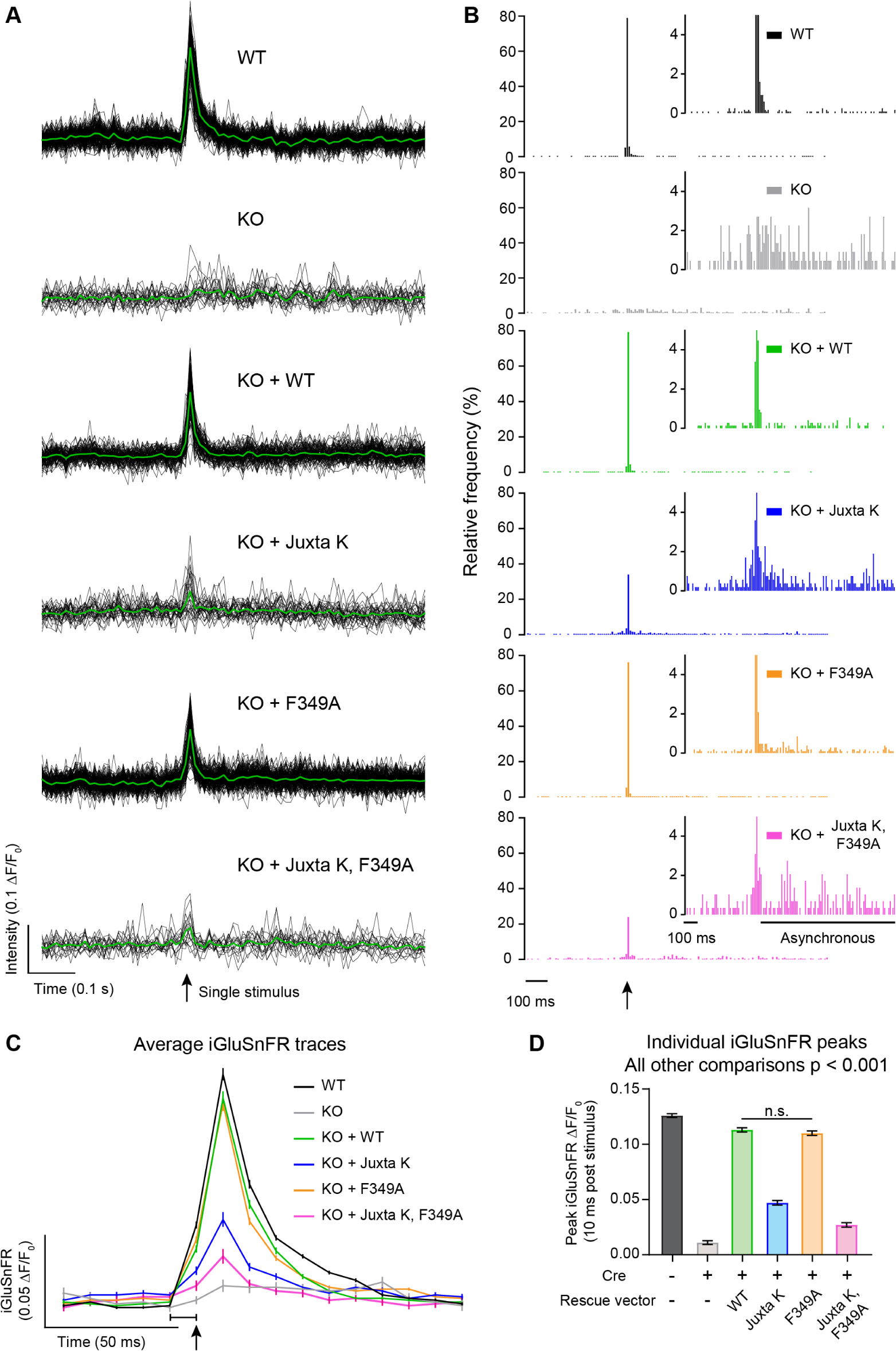
Lysine residues in the syt1 juxtamembrane linker regulate evoked neurotransmitter release. **A**) Representative iGluSnFR traces from one field of view after a single field stimulus (indicated by a black arrow) at 100 Hz. The plots show the iGluSnFR responses from multiple individual ROIs (black) and the average responses (green). Note that the KO, KO + Juxta K and KO + Juxta K, F349A conditions resulted in fewer responses after stimulation. **B**) Histograms of iGluSnFR (ΔF/F0) peaks plotted using 10 ms bins. The timing of the stimulus is indicated with a black arrow. Peaks were binned over the entire 1.5 s of recording. Conditions were color-coded as follows: WT (black), syt1 KO (grey), KO + WT (green), KO + Juxta K (blue), KO + F349A (orange), KO + Juxta K, F349A (magenta). The histograms include combined data from three independent trials. Y-axis zoom-in histograms of iGluSnFR (ΔF/F0) peaks shown as an inset to emphasize the presence of asynchronous release. **C**) Average traces of iGluSnFR ΔF/F0 from a single field stimulus, indicated by a black arrow, optically recorded at 100 Hz. Note, the 1 ms stimulus was applied in the last ms of the frame that is indicated by a black bracket; this causes a small portion of the rise in signal to be captured within that frame. Conditions are labeled in the figure and the same color scheme is used throughout. Briefly, WT (black), syt1 KO (grey), WT syt1 rescue (green), Juxta K mutant rescue (blue), F349A mutant rescue (orange), and the double Juxta K + F349A mutant rescue (magenta). Values are means with 95% CI error bars; the number of ROIs analyzed were: WT (1155), syt1 KO (212), WT rescue (748), Juxta K rescue (547), F349A rescue (1128), and double mutant rescue (324), collected from three separate trials. **D**) Average peak iGluSnFR responses 10 ms post stimulus from each condition. Quantification of data in panel (C). Values are mean +/- SEM from three independent trials. These data were normally distributed and only WT rescue and F349A mutant rescue were statistically similar, all other groups are different using ordinary ANOVA with Holm Sidak’s correction. All comparisons and statistical analysis are provided in Fig. 4-source data.

**Fig. 5.**
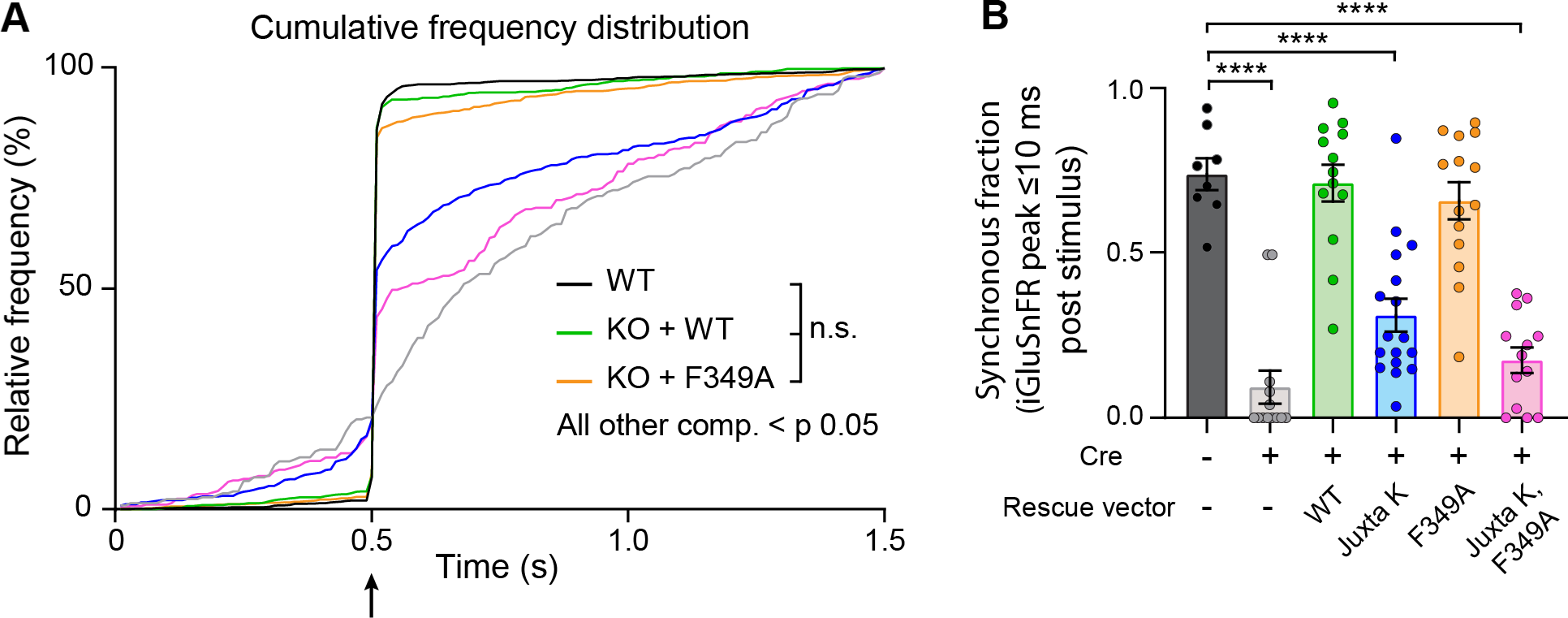
Lysine residues in the syt1 juxtamembrane linker regulate synchronized neurotransmitter release **A**) Cumulative frequency distribution of glutamate peaks throughout imaging, analyzed from Fig. 4; these data were not normally distributed. Using Kruskal-Wallis test with Dunn’s correction, no difference detected between WT, WT rescue, and F349A mutant rescue. All other comparisons were significantly different with p < 0.05. Full statistics are included in Fig. 5A-source data. **B**) Synchronous fraction of each condition quantified. Synchronous release defined as percentage of iGluSnFR ΔF/F0 peaks within 10 ms following a single stimulus (data are from Fig. 4 C and D) from an entire field of view. Values are mean +/- SEM from three independent trials. These data were normally distributed, with 8 to 17 field of views for each group; **** denotes p- value < 0.0001 between WT and labeled conditions using ANOVA with Holm-Sidak’s correction. All comparisons and statistical analysis, provided in Fig. 5B-source data.

### Syt1 self-association is required for clamping spontaneous release

Careful inspection of the iGluSnFR signals, before an action potential was delivered, indicated that some of the syt1 mutations also affected basal fusion rates (Fig. 5A). Indeed, it is well documented that under resting conditions, syt1 serves as a fusion clamp that inhibits spontaneous release (13-15, 43, 44). We therefore turned to electrophysiological analysis of the spontaneous (i.e. mini) fusion rates in syt1 KO neurons that expressed each construct (Fig. 6). We focused on inhibitory GABAergic minis (mIPSCs) as they are more reliant on syt1 than are glutamatergic minis (45) (Fig. 6). These experiments revealed yet another correlation: mutations that impair syt1 self-association also result in concomitant increases in spontaneous fusion rates (Fig. 6A and 6B). More specifically, F349A partially disrupted the ability of syt1 to clamp minis, as previously reported (37), while the Juxta K mutant had an even stronger effect; mutating both regions almost completely abolished the clamping activity of syt1 (Fig. 6A and 6B). The mini amplitudes were unchanged across all conditions (Fig. 6C).

**Fig. 6.**
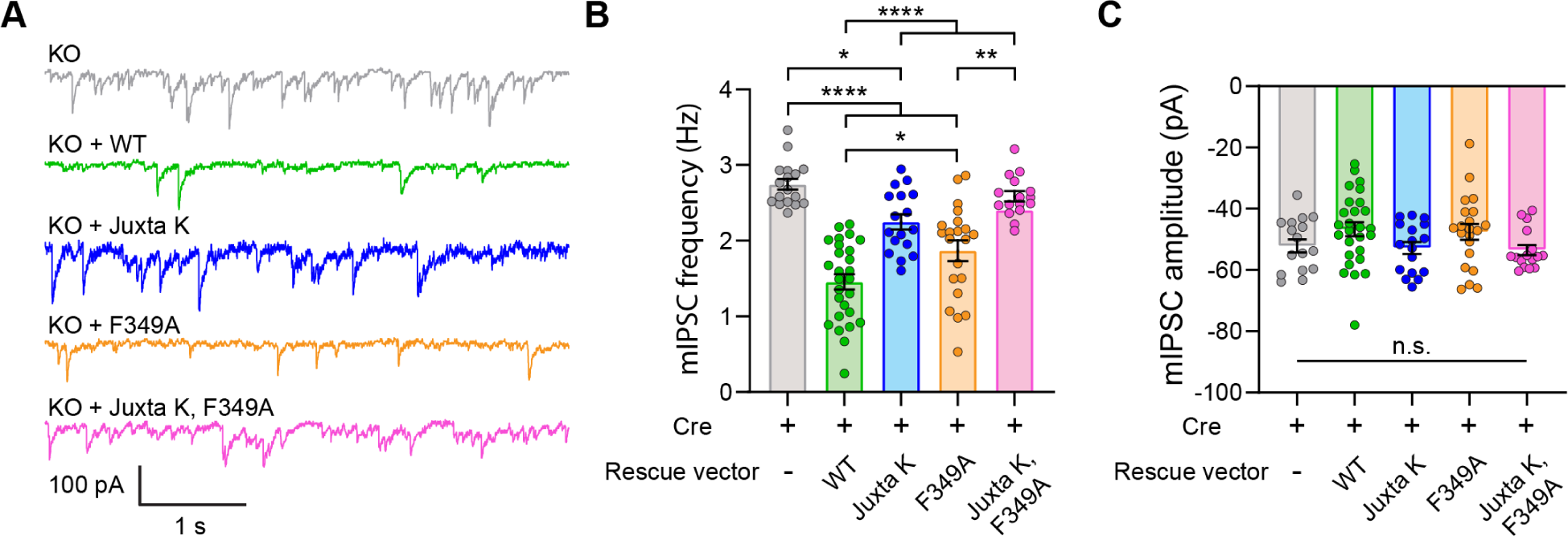
Lysine residues in the syt1 juxtamembrane linker regulate spontaneous neurotransmitter release. **A**) Representative traces of mIPSCs from syt1 KO neurons (grey) or syt1 KO expressing WT rescue (green), Juxta K mutant rescue (blue), F349 mutant rescue (orange), and Juxta K + F349A mutant rescue (magenta) syt1. **B**) Quantification of mIPSC frequencies from neurons expressing the various syt1 constructs. mIPSC frequency was 2.7 ± 0.07 Hz in syt1 KO neurons (mean ± SEM; n = 17), 1.5 ± 0.10 Hz in WT syt1 rescued cells (n = 27), 1.9 ± 0.14 Hz in F349A rescued cells (n = 21), 2.2 ± 0.10 Hz in Juxta K rescued cells (n = 17), and 2.4 ± 0.0.08 Hz in Juxta K, F349A rescued cells (n = 16). **C**) The quantification of mIPSC amplitudes after expression of the various syt1 constructs described in panel A. * denotes p-values < 0.05; ** denotes p-values < 0.01 and **** denotes p-value < 0.0001 determined by ANOVA using Tukey’s multiple comparisons test. All comparisons and statistical analyses are provided in Fig. 6-source data.

Together, the physiology experiments described in this study support a model in which syt1 must multimerize in order to clamp minis and to drive and synchronize evoked SV exocytosis.

## DISCUSSION

Cell-based experiments suggest that syt1 functions in the SV cycle as an oligomer (13, 22). Indeed, density gradient fractionation (24) and co-immunoprecipitation (46) experiments support the notion that syt1 oligomerizes in the presence of detergent. However, the stoichiometry and structure of these oligomers, as well as the effect of Ca^2+^ ions and phospholipids on self-association, remain unresolved issues. Hence, whether oligomerization impacts syt1 function remains unclear. It is therefore crucial to address the ability of syt1 to homo-multimerize on lipid bilayers, in the absence of detergent, under physiologically relevant conditions. In the current study, we first addressed this by conducting DLS, EM and AFM measurements in a reconstituted syt1•membrane systems, under detergent-free aqueous conditions. We observed that the complete cytoplasmic domain of syt1 does in fact form multimers on membranes under relatively native conditions. Importantly, anionic phospholipids are an essential cofactor for self-association (Fig. 1C and *SI Appendix*, Fig. S8).

We emphasize that our *in vitro* experiments utilized the complete cytoplasmic domain of syt1, residues 80-421, which includes the entire juxtamembrane linker between the TMD and the C2A domain. Residues 80-95 have largely been overlooked in previous studies examining syt1 function, but two studies suggested that this region might be important for function. In one report, a peptide corresponding to residues 80-98 of syt1 reduced neurotransmitter release, potentially by inhibiting syt1-membrane interactions (47). Also, as mentioned above, the juxtamembrane linker was reported to promote membrane binding and to mediate syt1 glycine zipper interactions (38). Strikingly, we did not observe rings or other multimeric structures when using a shorter fragment that began at residue 96 (unpublished observations); this fragment lacks the juxtamembrane lysine patch. These observations might seem to be at odds with the fact that rings on lipid monolayers were initially observed by EM with a syt1 C2AB domain that lacked the entire juxtamembrane linker (143–421) (32), but in that study, ring formation required low ionic strength buffers (5 - 15 mM KCl) and 40% PS. Indeed, syt1 self-association is highly dependent on the salt concentration of the buffer (29) and supraphysiological PS promotes calcium-independent membrane binding (48). Subsequent EM studies continued to include a low salt step during sample processing; however, it was later determined that the juxtamembrane linker could stabilize ring formation (36) after buffer exchange, from 5 mM to 100 mM KCl, prior to imaging. In short, both mutagenesis and truncation experiments demonstrate that the juxtamembrane linker is required for the cytoplasmic domain of syt1 to form homo-multimers at physiological ionic strength.

Under our aqueous conditions, syt1(80–421) formed large ring-like structures, with diameters ∼180 nm, on lipid bilayers; in contrast, previous reports of negative stain EM analysis revealed rings that were 17-45 nm in diameter (32). Another striking difference was that in our AFM experiments Ca^2+^ promoted self-assembly, while in the EM studies, Ca^2+^ dissolved the rings. Interestingly, Wang *et al*. (2017) also found that Ca^2+^ stabilized syt1 rings that had assembled in solution after binding a variety of polyanionic compounds. Differences in experimental conditions and sample handling steps, in EM versus AFM experiments, are likely to underlie the observed differences in multimeric structure and regulation. Specifically, AFM facilitates imaging of syt1 on lipid bilayers in aqueous buffer; EM yields greater resolution but involves extensive sample preparation and images are obtained on monolayers under vacuum. At present, we believe that the multimeric structures observed in our system are distinct from the previous reports of syt1 rings.

Interestingly, after neutralizing the juxtamembrane lysines, we observed small rings by AFM that were comparable to the size of the rings that were observed via EM. This may reflect the fact that the majority of the EM studies were, again, conducted using a truncated form of syt1 (143-421) that lacked the crucial juxtamembrane lysine-rich segment (28, 32, 35, 37). We went on to examine all three mutations (K326,327A (27, 34), F349A (28) and R398,399Q (29)) that have been implicated in oligomerization activity mediated by the C2B domain. None of these substitutions abolished self-association by themselves, but our AFM imaging found that each set of mutations completely disrupted the self-association in the Juxta K mutant background (Fig. 2C).

We therefore conclude that the juxtamembrane linker and the C2B domain of syt1 both contribute to a complex multimerization mechanism. The formation of large and small rings thus appears to involve somewhat distinct structural elements; the precise experimental conditions might determine which element dominates.

We took advantage of the mutations that impaired syt1 self-association *in vitro*, under relatively physiological aqueous conditions, and conducted functional assays in neurons. Deletion of the entire juxtamembrane linker has been shown to disrupt syt1 function, perhaps by altering the ability of the C2-domains to engage effectors (49). However, the role of the Juxta K region in SV exocytosis had not been previously explored via a mutagenesis approach. We first established that each mutant was efficiently targeted to SVs with proper topology, and then conducted physiology experiments that revealed a clear correlation between the ability of the mutations to disrupt multimerization activity with their ability to disrupt the clamping activity of syt1, resulting in higher rates of spontaneous SV release. Moreover, impairment of syt1 self-association was also correlated with reductions in peak glutamate release, and the desynchronization of this release, in response to single action potentials. Interestingly, a recent study using a reconstituted vesicle fusion assay proposed that syt1 must first clamp fusion in order to become subsequently responsive to Ca^2+^ (50). Together with our *in vitro* studies, these cell-based findings strongly suggest that syt1 must assemble into multimers, mainly via its juxtamembrane lysine-rich patch, but also with contributions from the C2B domain, in order to inhibit spontaneous release and to drive rapid and efficient evoked release.

We reiterate that large clusters observed by DLS and the ring-like structures that we observed by AFM are unlikely to form on SVs *in vivo*, as there are only ∼15 copies of syt1 per vesicle (23). In contrast, the syt1 copy number in our *in vitro* systems were largely unrestricted, and this is expected to exaggerate the oligomeric state. Still, these supraphysiological structures served as a platform to assay for regulatory factors (lipids, Ca^2+^), and structural elements, that govern syt1 self-association on lipid bilayers. Although the precise structural arrangement and oligomeric status of syt1 on SVs is still unknown, our data suggest that the ∼15 molecules of syt1 per SV would likely multimerize with one-another *in vivo*. We also note that ∼20-25% of neuronal syt1 resides in the plasma membrane of presynaptic boutons (51–53), and this pool could also potentially play a role in exocytosis by forming large (>15 copies) oligomers. Regardless, the *in vitro* experiments reported here demonstrate that highly purified syt1 self-associates under native conditions and these assays provided a means to monitor the disruption of this activity via mutagenesis to, in turn, guide functional experiments. Again, our pHluorin experiments largely rule-out a role for juxtamembrane-mediated oligomerization in endocytosis (Fig. 3F), but we demonstrate that syt1 self-association clearly plays a role in clamping spontaneous release (Fig. 6) and in determining the extent and synchronization of evoked SV exocytosis (Fig. 4 and 5).

At present, it is unclear as to how the oligomerization of syt1 contributes to its ability to regulate SV fusion. It seems likely that Juxta K-mediated multimerization of syt1 on the surface of SVs would, in addition to facilitating C2B-C2B interactions, influence how the C2-domains engage with binding partners on the pre-synaptic plasma membrane. For example, syt1 oligomerization might serve to ‘order’ SNARE proteins around the fusion pore, via direct physical interactions with t-SNAREs (10, 11, 39). Oligomerization could also play additional roles in the regulation of release by: adding mass to the fusion complex to drive pore dilation, orienting the C2-domains of syt1 to mediate its distinct effects on spontaneous and evoked release (15), allowing copies of syt1 to functionally cooperate with one another via direct physical interactions (22), or allowing groups of syt1 molecules to penetrate bilayers (29) and rearrange the phospholipids, as an ensemble, to drive fusion pore transitions. These issues will be addressed in future studies and will be facilitated by the robust AFM approach that, for the first time, allows the study of syt1 self-association under physiological conditions.

## METHODS

### Reagents

1,2-dioleoyl-sn-glycero-3-phosphocholine (PC), 1,2-dioleoyl-sn-glycero-3-phospho-l-serine (PS), 1,2-dioleoyl-sn-glycero-3-phosphoethanolamine (PE), 1-palmitoyl-2-oleoyl-sn-glycero-3- phospho-(1’-rac-glycerol) (PG) and 1,2-dioleoyl-sn-glycero-3-phospho-(1′-myo-inositol-4′,5′- bisphosphate) (PI(4, 5)P2) were obtained from Avanti Polar Lipids. HEPES was from Fisher Scientific, calcium chloride solution (1.0 M) was from Fluka Analytics, and all other chemicals were from Sigma-Aldrich.

### Recombinant proteins

Recombinant rat syt1 was purified from *E. coli* (BL21) as an N-terminally tagged his6-SUMO fusion protein. Protein expression was induced by the addition of 200 µM isopropyl β-d-1- thiogalactopyranoside (IPTG) when the OD600 of the culture reached 0.6 – 0.8. Bacterial pellets were lysed by sonication in 50 mM Tris, 300 mM NaCl, 5 % glycerol, 5 mM 2-mercaptoethanol, 1 % Triton X-100 (pH 7.4) plus a protease inhibitor cocktail (Roche). The samples were also incubated with RNase and DNase (10 µg/ml) to prevent nucleic acid mediated aggregation(54). Insoluble material was removed by centrifugation at 4000 rpm for 15 minutes, and the supernatant was incubated with Ni-NTA agarose, followed by washing the beads with 50 mM Tris, 1 M NaCl, 5 % glycerol (pH 7.4). Protein was liberated from the Ni-NTA agarose by overnight incubation with 0.5 µM recombinant SENP2 protease. A final FPLC purification was performed by running the samples through a Superdex 200 Increase 10/300 GL column in 25 mM HEPES, 300 mM NaCl, 5 % glycerol, 5 mM 2-mercaptoethanol (pH 7.4). Samples were subjected to SDS-PAGE and protein concentration was determined by staining with Coomassie blue, using BSA as a standard.

### DLS analysis

DLS of syt1(80–421) was performed using a DynaPro NanoStar Dynamic Light Scattering instrument (Wyatt Technology). The syt1(80–421) protein (2 µM) was suspended in 25 mM HEPES, 100 mM KCl, with and without 100 µM 1,2-dihexanoyl-sn-glycero-3-phospho-L-serine (6:0 PS) and average diameter distributions were determined. Each of the WT and mutant syt1(80–421) samples were analyzed in triplicate with consistent results. The DLS samples were then blinded and imaged by EM as previously described (55).

### AFM imaging

Thirty µl of liposomes (1mM stock solution; PC/PS/PIP2, 72:25:3 or PC/PS, 80:20, extruded with a 100 nm filter) were suspended in 270 µl imaging buffer (25 mM HEPES, pH 7.4, 150 mM potassium gluconate, 0.5 mM EGTA) with or without 1.5 mM CaCl2, and deposited onto freshly cleaved mica (20 mm diameter discs). After incubating for 30 min, the sample was rinsed with 150 µl of the same buffer six times, using a pipette, to remove unabsorbed material; substrate was covered by buffer at all times. Purified syt1(80–421) was suspended in the same buffer and added to the sample. After a 20-min or 6-hour incubation period, the sample was again rinsed repeatedly to remove unbound protein, and the final sample volume was adjusted to 300 µl. AFM imaging was carried out with a Agilent 5500 Scanning Probe Microscope in AAC mode with silicon nitride probes (FastScan-D, Bruker or BL-AC40TS, Oxford Instruments). Images were captured with minimum imaging force at a scan rate of 2 Hz with 512 lines per area. Data analysis was performed with PicoImage 5.1.1. Particle volume was determined using the peak/dip volume tool (within the PicoImage software) line-by-line. Ring structure diameter was defined as the distance between the two highest points on each side of a cross-section across the imaged structure. Protein coverage on lipid bilayers were determined by setting the height threshold to anything one nm or more above the bilayer surface.

### Giant unilamellar vesicle and supported lipid bilayer microscopy

Giant unilamellar vesicles (GUVs) and rhodamine-PE labeled supported lipid bilayers (SLB) were imaged using a Zeiss 880 Airyscan microscope. GUVs were prepared by drying 15 µl of 1 mM DOPC/DOPS lipids, plus 0.1% rhodamine-PE in three locations onto indium tin oxide coated glass slides and then performing electroformation with 10 Hz, 4 V alternating current for two hours in 1 mM HEPES, 200 mM Sucrose solution. The buffer also contained 10 µM Alexa-647 dye. The excess dye was removed by filtration. The SLBs were prepared on mica discs as described above, except with the inclusion of 0.1% rhodamine-PE within the lipid mixture.

### Cell Culture

Syt1 floxed neurons were prepared as previously described (14). Briefly, hippocampal neurons were dissected at P0 from syt1 floxed mouse strain (Quadros et al., 2017), trypsinized (Corning; 25-053-CI), triturated, and plated on glass coverslips (Warner instruments; 64-0734 (CS-18R17)) coated with poly-D-lysine (Thermofisher; ICN10269491) and EHS laminin (Thermofisher; 23017015). Neurons were grown for at least 14 days in Neurobasal-A (Thermofisher; 10888-022) medium supplemented with B-27 (2% Thermofisher; 17504001), Glutamax (2 mM Gibco; 35050061), and pen/strep before experiments. For virus preparation, HEK293T cells (ATCC) were cultured following ATCC guidelines and were tested for mycoplasma contamination using the Universal Mycoplasma Detection Kit (ATCC; 30-1012K), and validated as HEK293T cells using Short Tandem Repeat profiling by ATCC (ATCC; 135-XV) within the last year.

### Lentivirus production and use

Lentivirus production was performed as described previously (14). Syt1 expressing lentiviral constructs were subcloned into the FUGW transfer plasmid (FUGW was a gift from David Baltimore (Addgene plasmid # 14883) (56). Our lab has previously modified this construct, replacing the ubiquitin promoter with the human synapsin I promoter (57). Lentivirus expressing Cre was added to neuronal cultures at day one in-vitro (DIV), iGluSnFR and pHluorin constructs were also added at 1 DIV. Syt1 constructs were added at 5 DIV.

### Plasmid construction for lentiviral expression

All plasmids, unless otherwise noted, were constructed using our lab’s modified lentivirus backbone of choice derived from FUGW. The glutamate sensor is the same as used previously (14). The pHluorin construct used here was subcloned into our modified FUGW transfer vector from the original vGlut1-pHluorin construct (58). All synaptotagmin 1 constructs were subcloned into our modified FUGW transfer plasmid from their original bacterial expression plasmids. For Cre expression, we used the transfer plasmid pLenti-hSynapsin-CRE-WPRE (pLenti-hSynapsin-CRE-WPRE was a gift from Fan Wang (Addgene plasmid # 86641) (59).

### Immunoblot analysis

Immunoblots were performed as described previously (14). Primary antibodies were: anti-syt1 (1:1000, 48) (lab stock; mAB 48; RRID:AB_2199314). Secondary antibodies were: goat anti-mouse IgG2b-HRP (Biorad, M32407; RRID:AB_2536647).

### Immunocytochemistry (ICC)

Immunocytochemistry was performed as previously described (14). Primary antibodies were: anti- syt1 (1:100, 48) (lab stock; mAB 48; RRID:AB_2199314) and anti-synaptophysin (SYP) (1:500) (SySy; 101 004; RRID:AB_1210382). Secondary antibodies used were: goat anti-guinea pig IgG-Alexa Fluor 546 (1:500) (Thermofisher; A-11074, RRID:AB_2534118) and goat anti-mouse IgG2b-Alexa Fluor 647 (1:500) (Thermofisher; A-21242; RRID:AB_2535811). Images in Fig. 3 were acquired on a Zeiss LSM 880 with a 63x 1.4 NA oil immersion objective using the Airyscan super-resolution detector. The same laser and gain settings were used for each condition. Images were deconvolved using automatic Airyscan settings and the same linear brightness and contrast adjustments were applied to all images for figure presentation.

### iGluSnFR imaging and quantification

iGluSnFR imaging and quantification was performed as previously described (14), with the following modifications. For single stimuli imaging, 150 frames were collected at 10 ms exposure (1.5 sec total) and a single field stimulus was triggered at half a second after the initial frame.

### pHluorin imaging

Live cell fluorescent imaging of pHluorin expressing neurons was carried out under the same conditions as iGluSnFR imaging. Briefly, images were acquired on an Olympus IX83 inverted microscope equipped with a cellTIRF 4Line excitation system using an Olympus 60x/1.49 Apo N objective and an Orca Flash4.0 CMOS camera (Hamamatsu Photonics). This microscope runs Metamorph software with Olympus 7.8.6.0 acquisition software from Molecular Devices. Imaging media was extracellular fluid (ECF), with 2 mM CaCl2. Single image planes were acquired with 500 ms exposure using a white OLED with standard GFP filters. Images were collected once a second for 3 minutes. A stimulation train was started 9 seconds into imaging. The train (200 stimuli in 10 seconds (20 Hz)) were triggered by a Grass SD9 stimulator through platinum parallel wires attached to a field stimulation chamber (Warner Instruments; RC-49MFSH). All biosensor imaging experiments were performed at 32-34 °C. Environment was controlled by a Tokai incubation controller and chamber.

### Colocalization quantification

Colocalization was measured as described previously (14), using Fiji for ImageJ and Just Another Colocalization Plugin (JACoP) (60).

### Electrophysiology

Miniature inhibitory postsynaptic currents (mIPSCs) were recorded using a Multiclamp 700B amplifier (Molecular Devices) and analyzed as previously described (45). Briefly, syt1 KO hippocampal neurons expressing WT, Juxta K, F349A or Juxta K + F349A at DIV 14–19 were transferred to a recording chamber with a bath solution containing the following (in mM): 128 NaCl, 5 KCl, 2 CaCl2, 1 MgCl2, 30 D-glucose, 25 HEPES, and 1 μM TTX, pH 7.3 (305 mOsm). Borosilicate glass pipettes (Sutter Instruments) were pulled by a Dual-stage glass micropipette puller (Narishige) and filled with an internal solution contained (in mM): 130 KCl, 1 EGTA, 10 HEPES, 2 ATP, 0.3 GTP, 5 QX-314 (Abcam), and 5 sodium phosphocreatine, pH 7.35 (295 mOsm). mIPSCs were pharmacologically isolated by bath applying D-AP5 (50 µM, Abcam) and CNQX (20 µM, Abcam) and acquired using a Digidata 1440B analog-to-digital converters (Molecular Devices) and Clampex 10 software (Molecular Devices) at 10 kHz. Neurons were held at −70 mV. All cells were equilibrated for ∼1 min after break-in before recordings started. Series resistance was compensated, and traces were discarded if the access resistance exceeded 15 ΜΩ for the entire duration. The collected miniature events were detected in Clampfit 11.1 (Molecular Devices) using a template matching search.

### Statistics

Values from analysis and number of trials (n) for each experiment are listed in the figure legends. All analysis was done using GraphPad Prism 7.04 (GraphPad Software Inc).

## ACKNOWLEDGMENTS

We thank the members of the Chapman Lab for providing helpful discussions, Julie Morasch for assistance with AFM experiments, and Noah D. Miller for assistance with plasmid and lentivirus preparation. This work was funded by the National Institutes of Health (grants MH061876 and NS097362 to E.R.C.). E.R.C. is an Investigator of the Howard Hughes Medical Institute.

## COMPETING FINANCIAL INTERESTS

The authors declare no competing financial interests.

## Notes

### Competing Interest Statement

The authors have declared no competing interest.

