## Supplemental figures for "Synaptotagmin 1 oligomerization via the juxtamembrane linker regulates spontaneous and evoked neurotransmitter release"

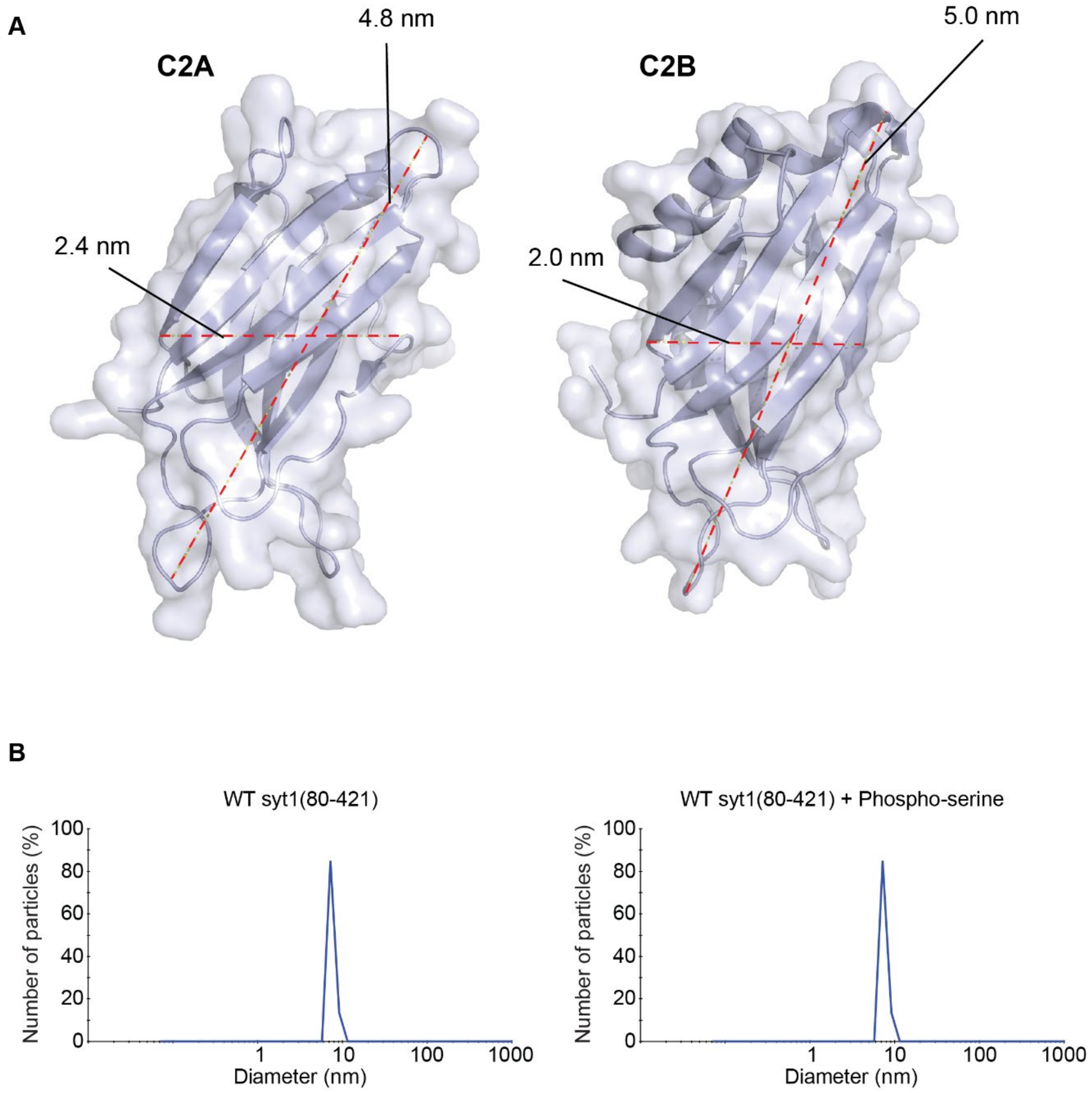


**Fig. S1.** Dynamic light scattering of the syt1 soluble domain in solution. **A**) Structures of syt1 C2A and C2B domains, with measured dimensions, derived from RCSB PBD files 5t0r and 2yoa, respectively. **B)** Representative dynamic light scatter results of 2 µM WT syt1(80-421) with and without the addition of 100 µM phospho-serine.

**A**

**Recombinant syt1(80-421) juxtamembrane sequences**

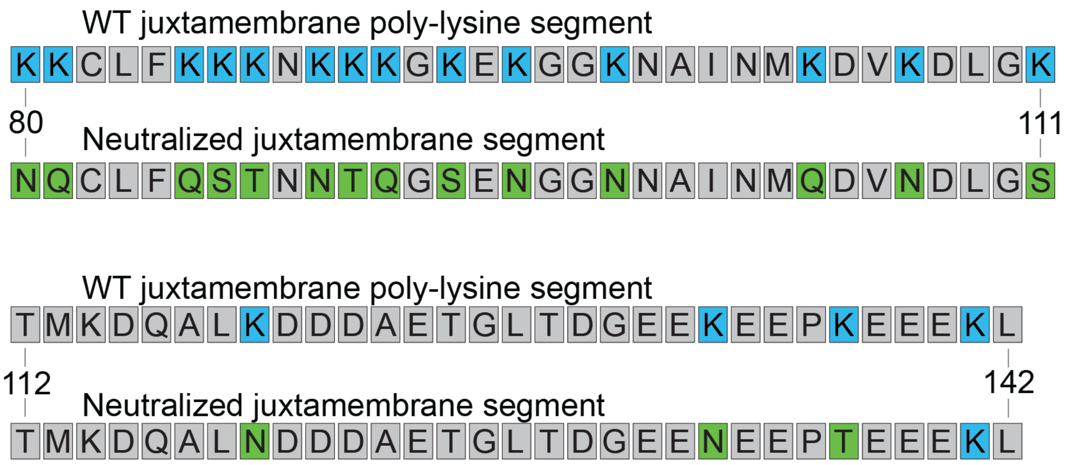


**B**

**Full length syt1 lentiviral expression juxtamembrane sequences**

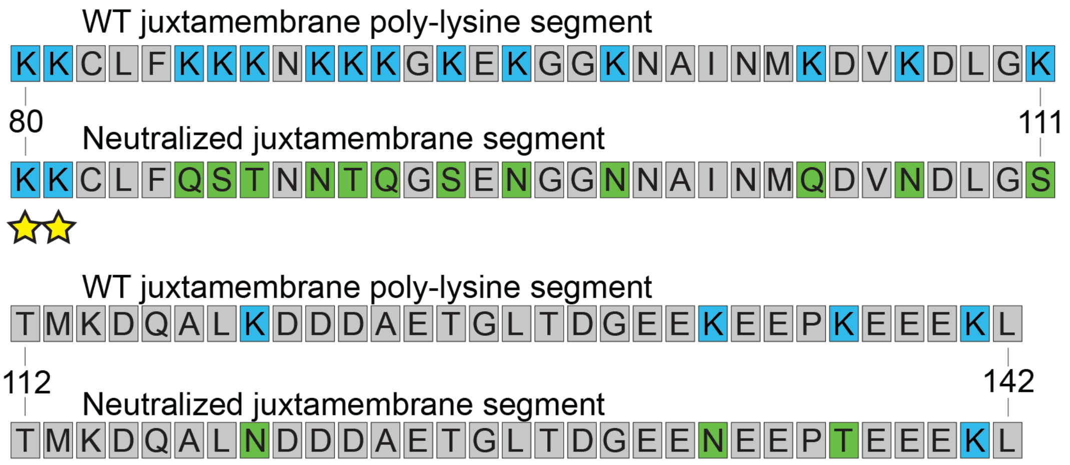


**Fig. S2.** Complete juxtamembrane amino acid sequences of WT and the Juxta K mutant form of syt1. **A**) For *in vitro* experiments, syt1(80-421) was used. The juxtamembrane sequences of WT and the Juxta K mutant are shown. WT lysine residues are in blue and neutralizing substitutions in the Juxta K mutant are shown in green. **B**) For cell-based experiments of WT and mutant forms of syt1, the full-length protein (residues 1-421) was used. In this case, two lysine resides, 80 and 81, were preserved to maintain proper topogenesis (41); these are marked with yellow stars. Color-coding is the same as in *panel (A)*.

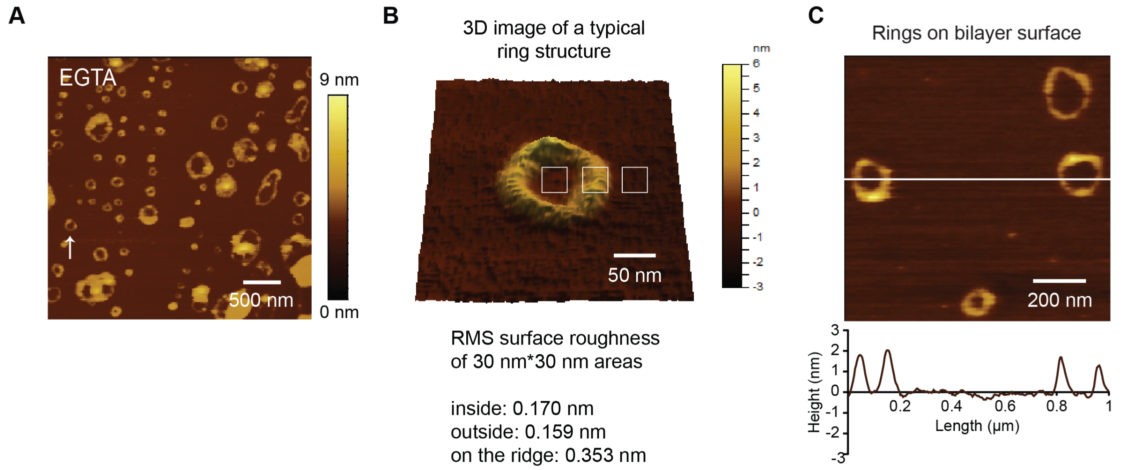


**Fig. S3.** Syt1(80-421) forms ring-like multimeric structures on the surface of lipid bilayers. **A**) Atomic force microscopy (AFM) topographical images of the entire cytoplasmic domain of syt1 (syt1(80-421)) on a supported lipid bilayer (72% DOPC, 25% DOPS, 3% $PIP_2$), under aqueous conditions, after a 6-hour incubation. Images were acquired at 9.8 nm/px. A white arrow indicates a representative oligomeric structure with a corresponding zoomed-in three-dimensional representation shown in *panel B*. **B**) Representative 3D image of a ring-like syt1 structure, derived from *panel A*, with surface roughness analysis. Root mean squared (RMS) surface roughness inside (0.170 nm) and outside (0.159 nm) the ring was compared to the roughness of the proteinaceous ring (0.353). **C**) Representative AFM image with topographical line scans across the syt1(80-421) structure that assembled on the surface of the supported lipid bilayer. Line scan shows the height of the internal surface matches the height of the surrounding bilayer (normalized to zero).

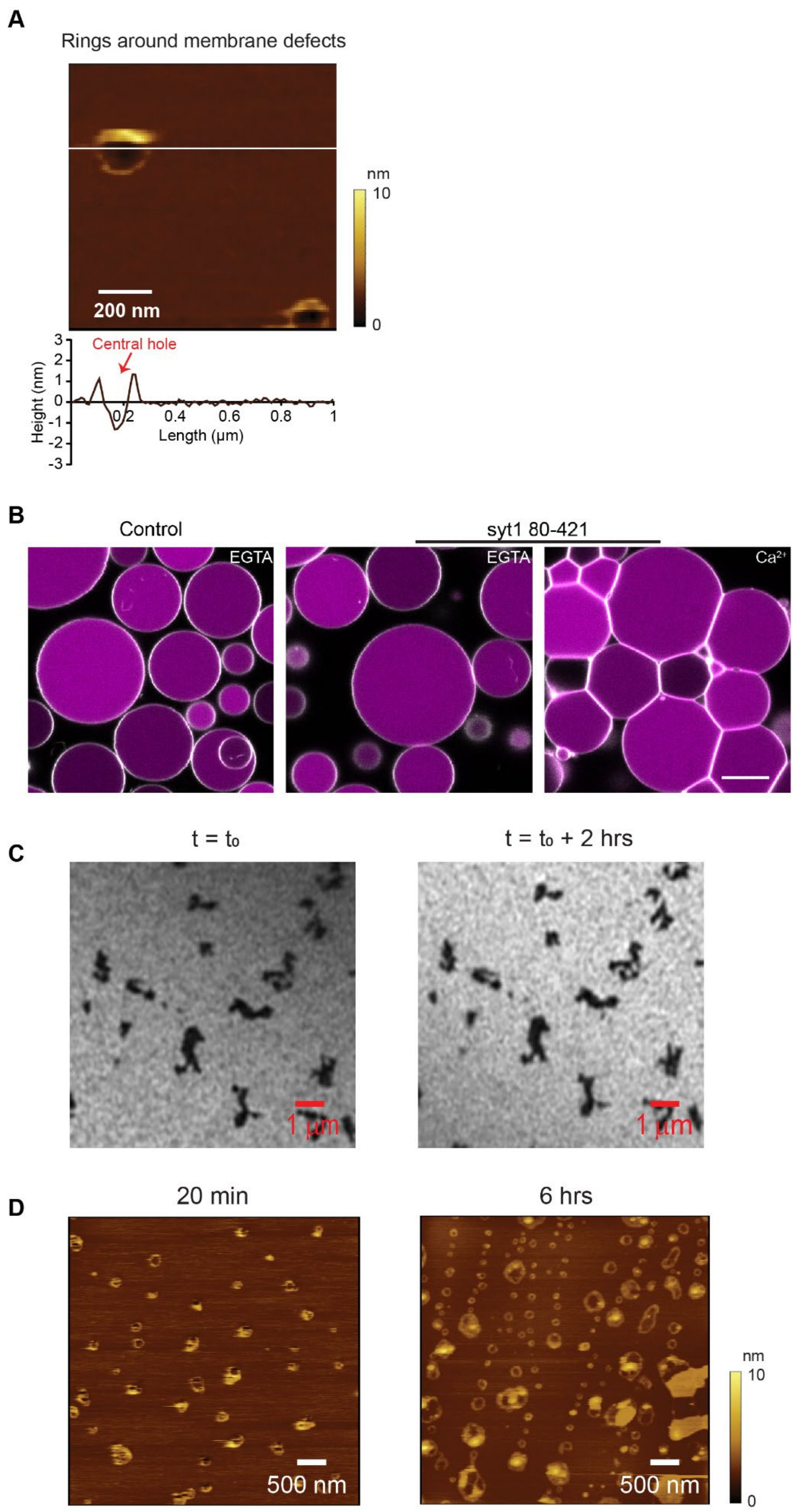
A
Rings around membrane defects
nm
10
0
200 nm
Central hole
Height (nm)
Length (µm)
B
Control
syt1 80-421
EGTA
EGTA
Ca²⁺
C
t = t₀
t = t₀ + 2 hrs
1 µm
1 µm
D
20 min
6 hrs
500 nm
500 nm
nm
10
0

**Fig. S4.** Ring-like syt1 structures around bilayer defects were excluded from analysis. **A**) Representative AFM image with topographical line scans across a syt1(80-421) structure that assembled around a membrane defect. All rings with a central hole were similarly identified by line scanning and were excluded from analysis. **B**) Confocal fluorescence microscopy of giant unilamellar vesicles (labeled with 0.1% Rhodamine-PE, white) composed of DOPC/DOPS (80:20) with encapsulated Alexa-647 dye (magenta). Samples were incubated for 20 minutes with 1 μM syt1(80-421) in 0.5 mM EGTA or 1 mM free $Ca^{2+}$. Note that after the addition of $Ca^{2+}$, the vesicles became clustered, due to syt1 bridging between vesicles. Scale bar represents 10 μm. **C**) PC/PS/PIP2 bilayers, containing 0.1% Rhodamine-PE were assembled on freshly cleaved mica discs and imaged by fluorescence microscopy. The stability of defects in the bilayer were assessed by imaging the same region twice, two hours apart. **D**) Representative AFM images of supported lipid bilayers that were incubated with 1 μM syt1(80-421) for twenty minutes or six hours. Note that these images were acquired from independent samples, after syt1(80-421) had been washed away. We found that AFM imaging in the presence of syt1(80-421) greatly affected the AFM tip stability, precluding the ability to visualize the real-time assembly of multimeric structures.

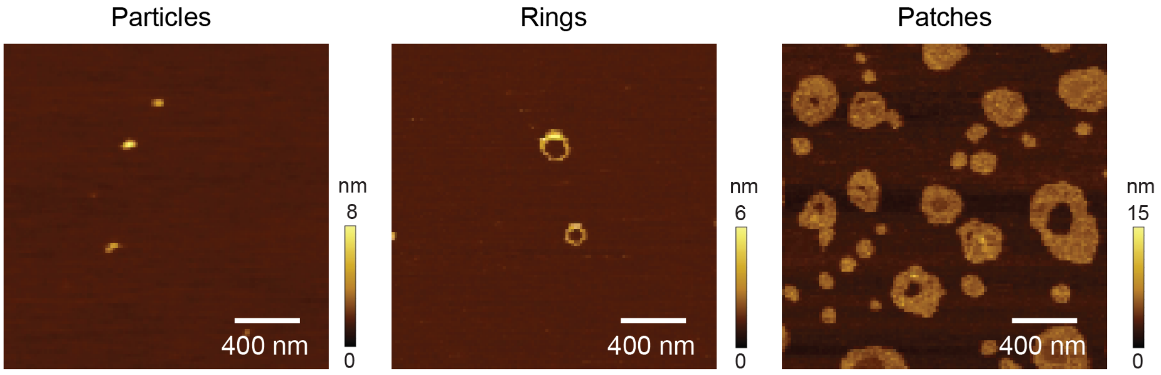


**Fig. S5.** The morphology of syt1(80-421) structures that assemble on supported lipid bilayers is concentration dependent. **A**) Representative AFM topographical images of syt1(80-421) forming particles (50 nM), rings (1 μM) or patches (3 μM) on supported lipid bilayers.

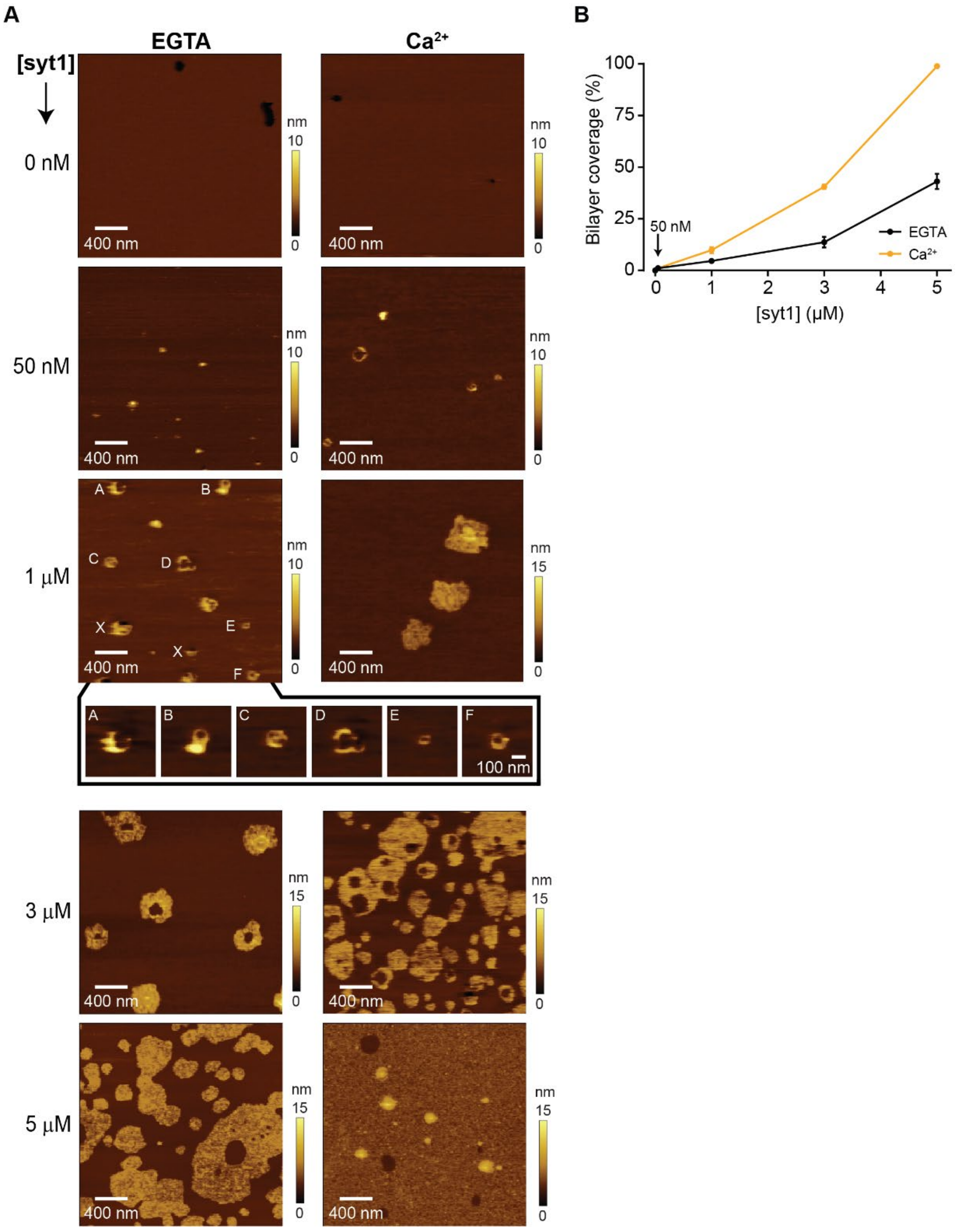


**Fig. S6.** AFM images of supported lipid bilayers as a function of syt1(80-421) protein concentration. **A**) Representative AFM topographical images of syt1(80-421), at the indicated concentrations, on supported lipid bilayers (72% DOPC, 25% DOPS, 3% $PIP_2$) in the presence (1 mM) or absence of $Ca^{2+}$ (0.5 mM EGTA). The images were acquired with a resolution of 9.8 nm/px. The inset under the 1μM syt1(80-421) condition in EGTA shows zoomed-in examples, with corresponding annotations, of typical structures. The white X symbols indicate protein structures surrounding lipid defects, which were excluded from our analysis. **B**) Protein coverage on lipid bilayers, at the indicated [protein], was calculated by setting the height threshold to ≥1 nm above the lipid bilayer surface. The error bars are standard deviation of protein coverage from three different scan areas in the same experiments. **C**) Representative histograms of dynamic light scattering size distributions of syt1(80-421) in solution in the presence or absence (0.5 mM EGTA) of 1 mM free $Ca^{2+}$ and/or 1,2-dihexanoyl-sn-glycero-3-phospho-L-serine (6:0 PS). Note that soluble 6:0 PS alone did not generate a measurable light scatter signal. The experiment was repeated 3 times with consistent results.

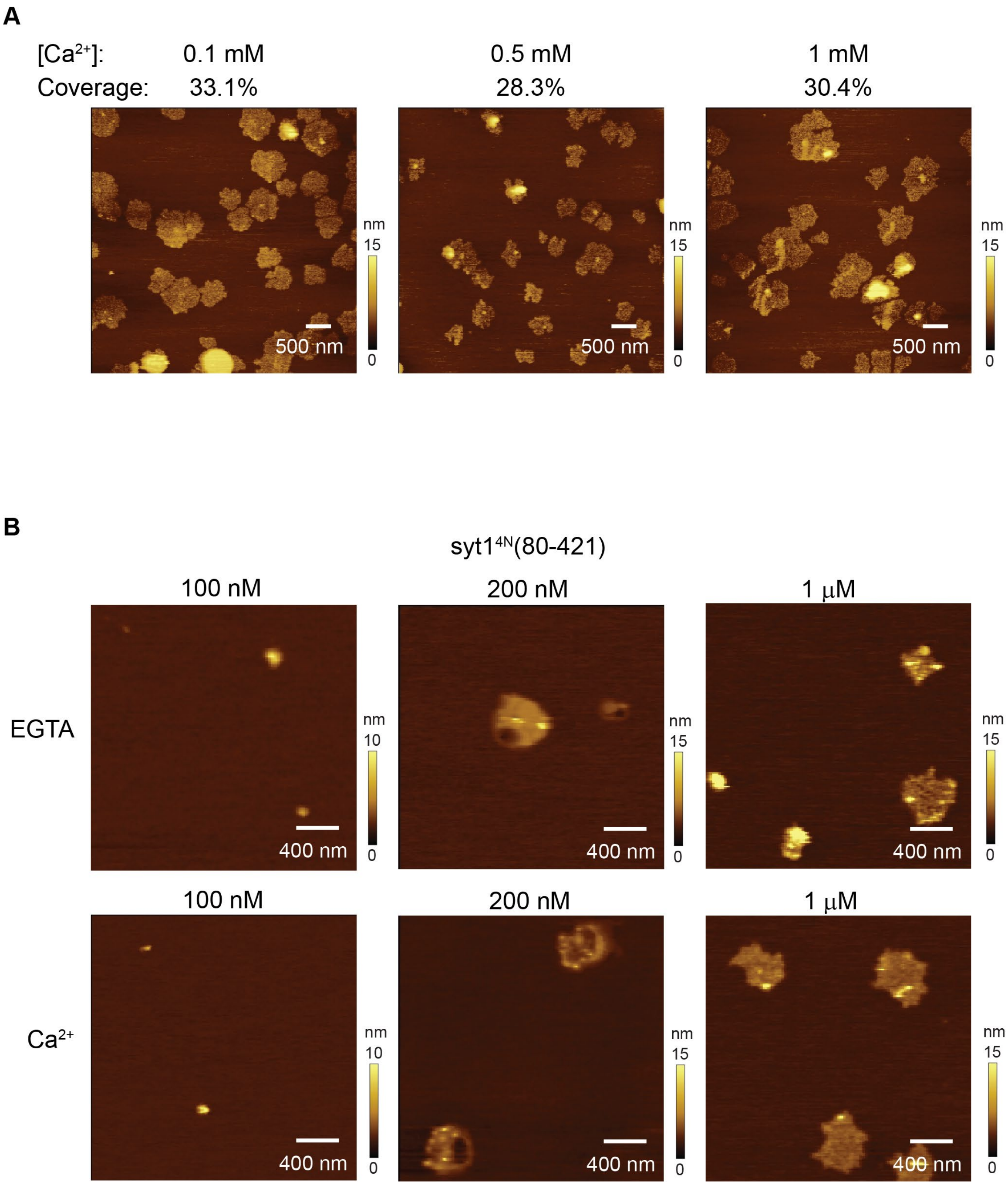


**Fig. S7.** $Ca^{2+}$ enhances syt1(80-421) self-association. **A**) Syt1(80-421) (1 μM) multimers remained fully formed as patches when the $[Ca^{2+}]$ was reduced from 1 to 0.5 and 0.1 mM. **B**) Substitution of acidic $Ca^{2+}$ ligands in both C2-domains of syt1 prevents $Ca^{2+}$-induced enhancement of syt1(80-421) oligomerization.

Representative AFM imaging of syt1$^{4N}$(80-421) (D230,232,363,365N) oligomerization on supported lipids bilayers in 0.5 mM EGTA or 1 mM $Ca^{2+}$.

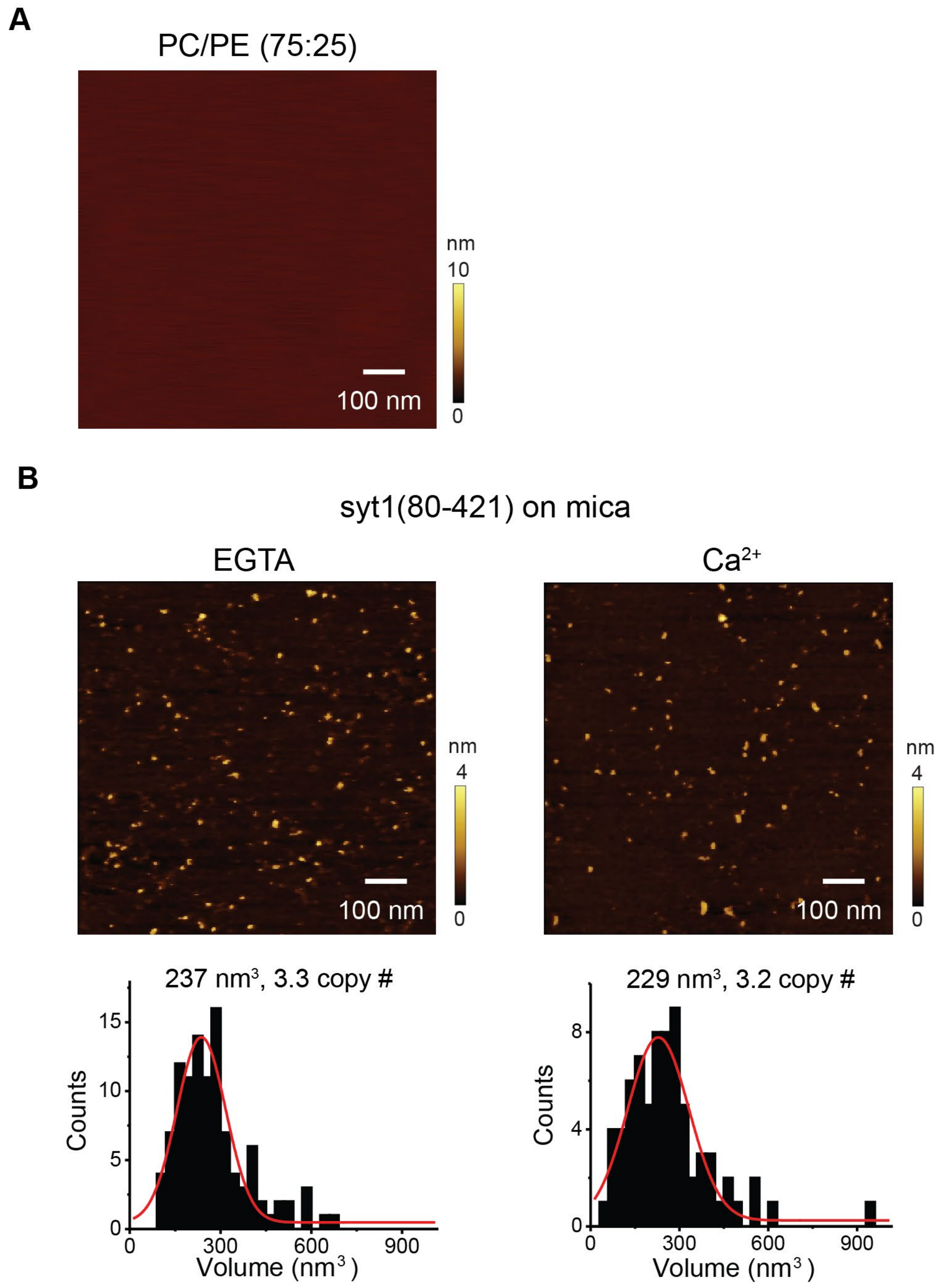


**Fig. S8.** Syt1 oligomerization on supported lipids bilayers requires anionic phospholipids. **A**) Representative AFM imaging of syt1(80-421) on a supported lipid bilayer composed of PC/PE (75:25). The anionic phospholipids, PS and $PIP_2$ were substituted with PE to enable efficient rupturing of large unilamellar vesicles on mica to generate the supported lipid bilayers. **B**) AFM imaging of syt1(80-421)

in the absence of supported lipid bilayer (bare mica) in 0.5 mM EGTA or 1 mM $Ca^{2+}$. Syt1(80-421) size distribution histograms for 0.5 mM and 1 mM $Ca^{2+}$ are shown below.

WT syt1(80-421) oligomeric structures on lipid bilayers

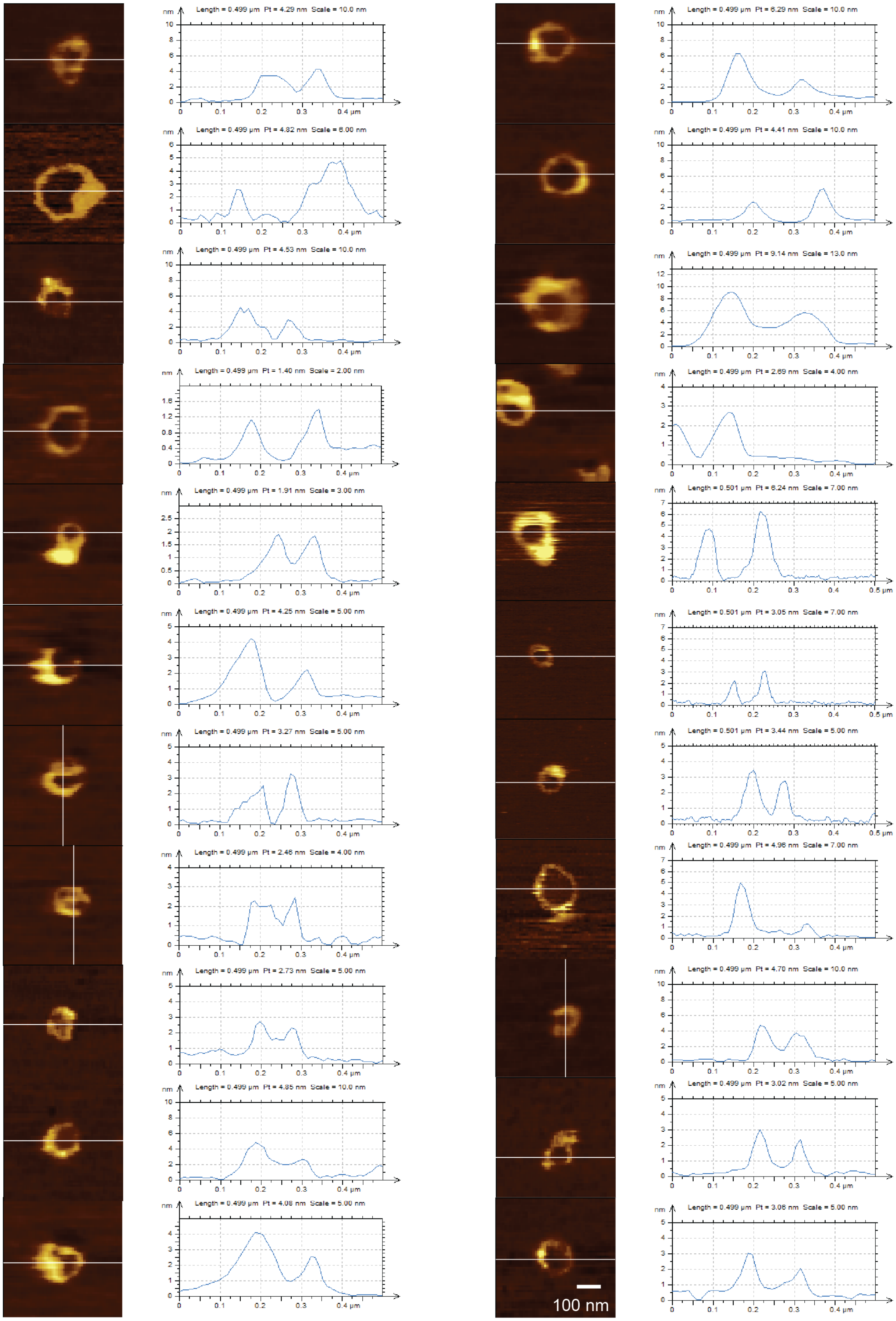

**Fig. S9.** WT syt1(80-421) forms ring-like structures on supported lipid bilayers in EGTA. Lateral height profiles confirm that only structures lacking a central hole were included in the analysis plotted in Fig. 2D.

K326,327A oligomeric structures on lipid bilayers

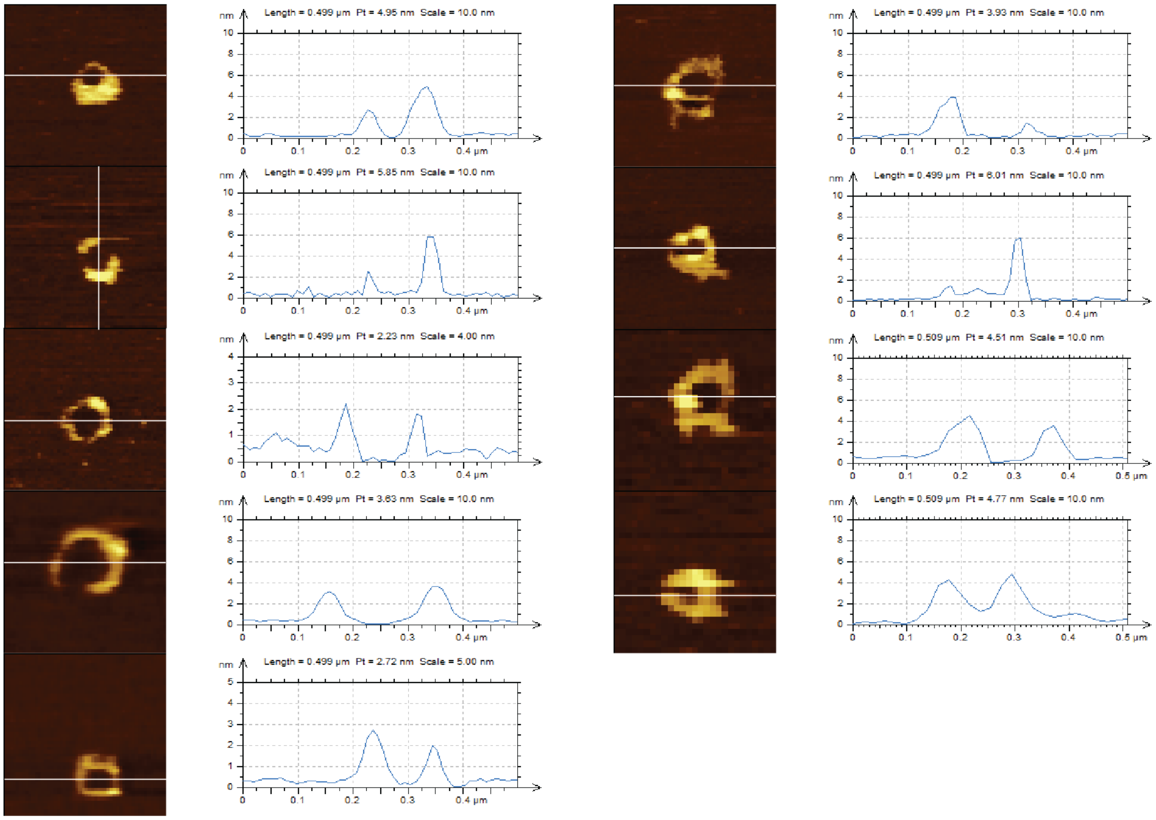


**Fig. S10.** The K326,327A syt1(80-421) mutant forms ring-like structures on supported lipid bilayers in EGTA. Lateral height profiles confirm that only structures lacking a central hole were included in the analysis plotted in Fig. 2D.

F349A oligomeric structures on lipid bilayers

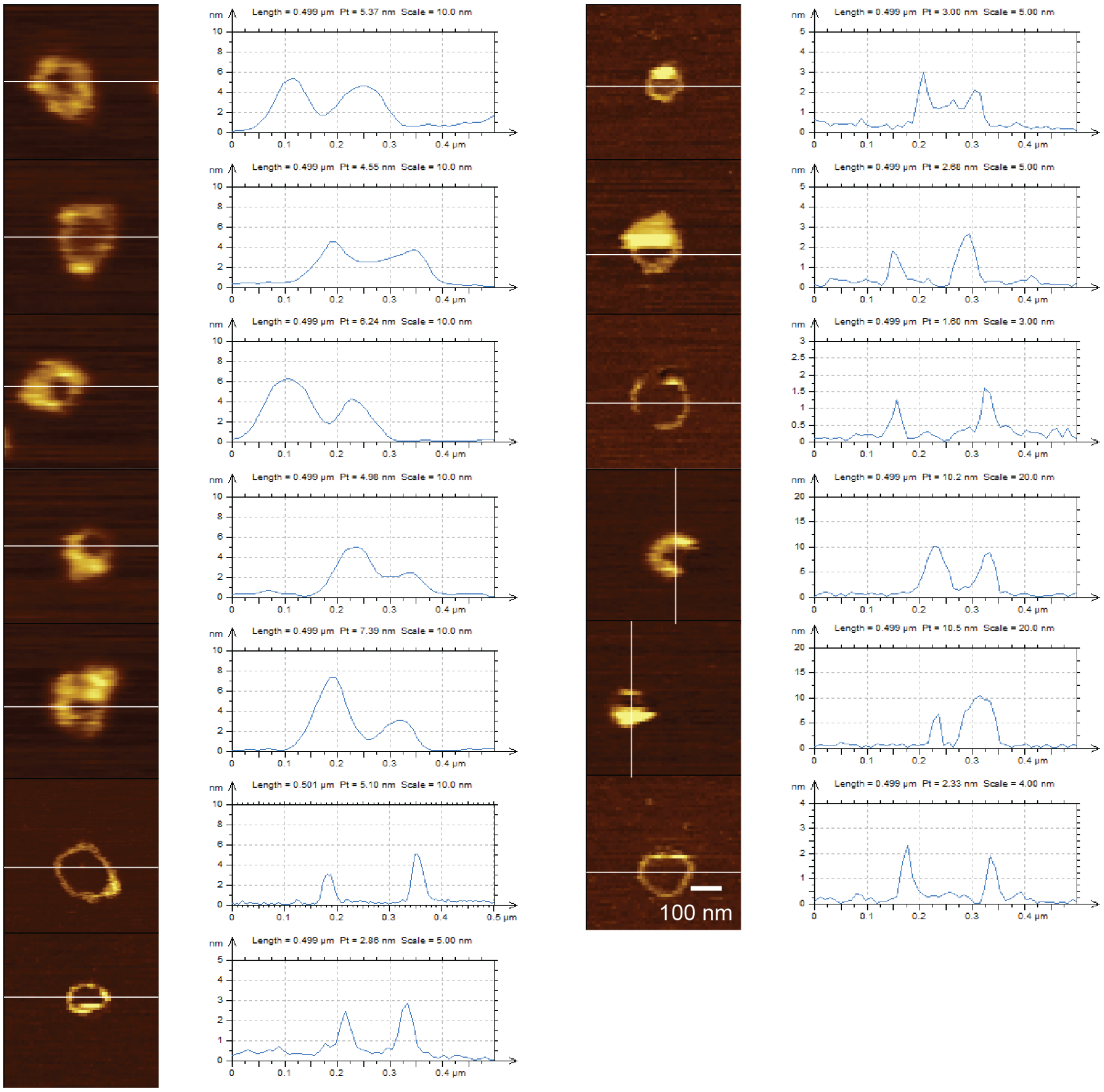


**Fig. S11.** The F349A syt1(80-421) mutant forms ring-like structures on supported lipid bilayers in EGTA. Lateral height profiles confirm that only structures lacking a central hole were included in the analysis plotted in Fig. 2D.

R398,399Q oligomeric structures on lipid bilayers

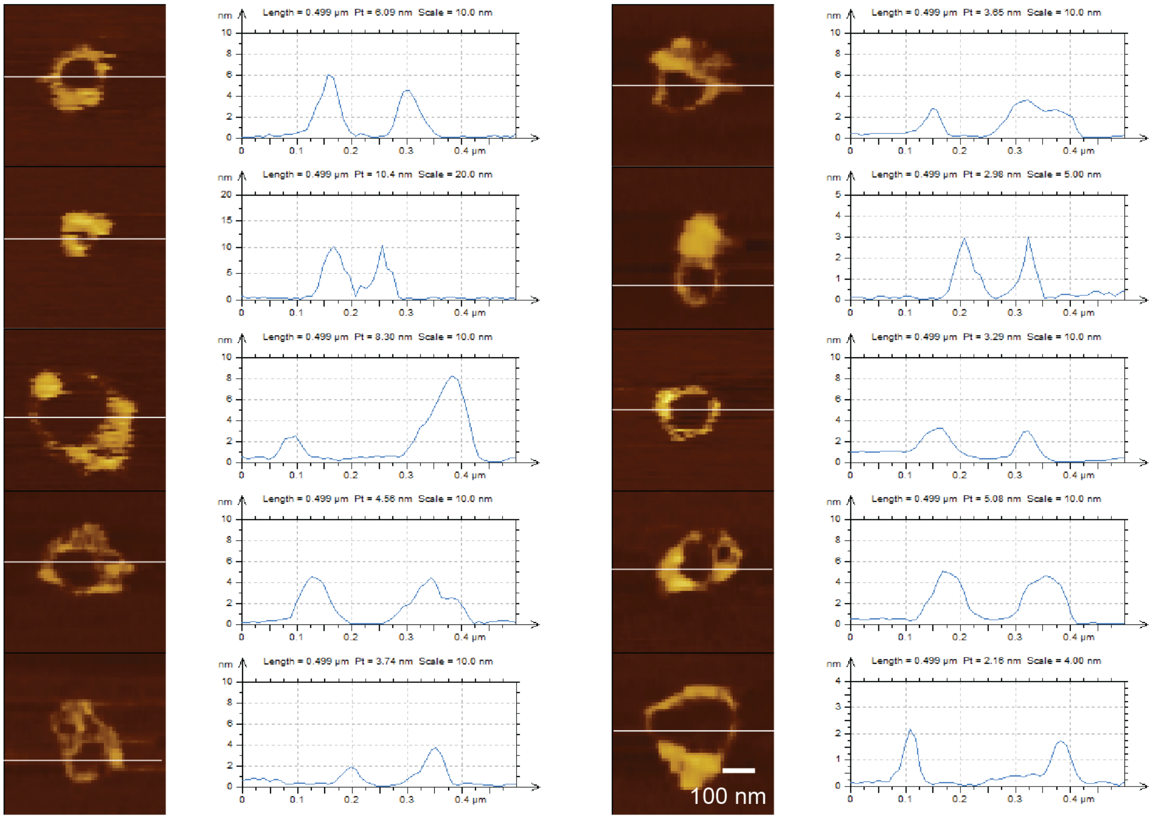


**Fig. S12.** The R398,399Q syt1(80-421) mutant forms ring-like structures on supported lipid bilayers in EGTA. Lateral height profiles confirm that only structures lacking a central hole were included in the analysis plotted in Fig. 2D.

Juxta K oligomeric structures on lipid bilayers

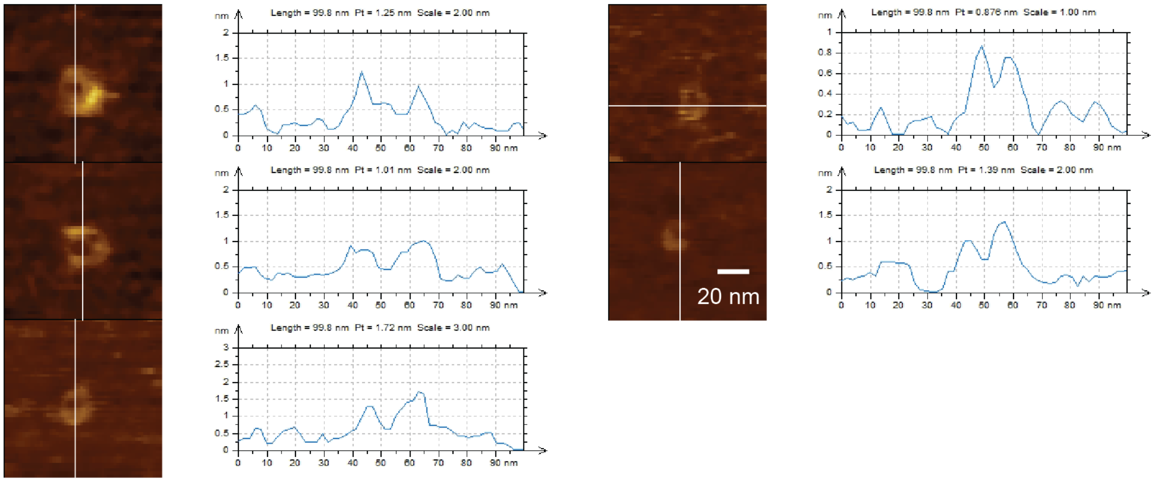


**Fig. S13.** The Juxta K syt1(80-421) mutant forms ring-like structures on supported lipid bilayers in 1 mM $Ca^{2+}$. No ring-like structures were observed in EGTA. Lateral height profiles confirm that only structures lacking a central hole were included in the analysis plotted in Fig. 2D.

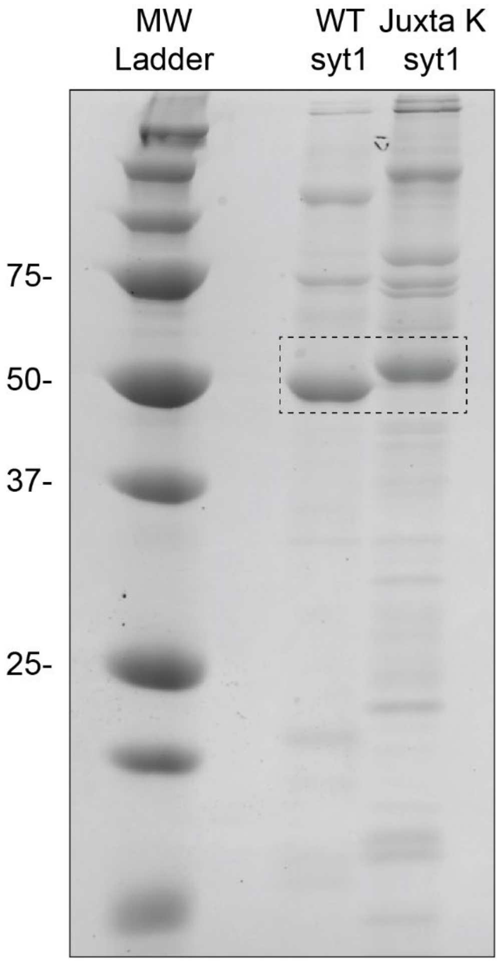


**Fig. S14.** Recombinant Juxta K mutant syt1 migrates through SDS-PAGE gels slower than WT syt1. Coomassie blue stained SDS-PAGE gel containing a molecular (MW) ladder and recombinant WT or Juxta K mutant syt1 that was purified from *E. coli*. The dotted box indicates the location of the recombinant syt1 protein.

**Table S1.** Statistics on syt1(80-421) morphology at increasing concentration with and without $Ca^{2+}$.

| Syt1(80-421) concentration (in 0.5 mM EGTA) | 30 nM | 50 nM | 200 nM | 1 mM | 3 mM |
|---|---|---|---|---|---|
| Proportion of particles (%) | 100 | 100 | 43.3 | 12.8 | 0 |
| Proportion of rings (%) | 0 | 0 | 23.3 | 78.7 | 0 |
| Proportion of Patches (%) | 0 | 0 | 33.3 | 8.5 | 100 |
| Ring size (nm) | NA | NA | 131±32 | 175±40 | NA |

| Syt1(80-421) concentration (in 1 mM free $Ca^{2+}$) | 30 nM | 50 nM | 100 nM | 1 mM | 3 mM |
|---|---|---|---|---|---|
| Proportion of particles (%) | 40 | 12.5 | 22.2 | 0 | 0 |
| Proportion of rings (%) | 60 | 62.5 | 55.6 | 0 | 0 |
| Proportion of Patches (%) | 0 | 25 | 22.2 | 100 | 100 |
| Ring size (nm) | 147±23 | 150±31 | 148±34 | NA | NA |

**Table S2.** Numerical values of individual iGluSnFR peaks associated with Fig. 4D.

| Condition | Average | SD | N | SEM | Change relative to WT |
|---|---|---|---|---|---|
| WT | 0.126 | 0.059 | 1155 | 0.001736 | 0.00 +/- 1.9% |
| KO | 0.011 | 0.025 | 212 | 0.001717 | -91.30 +/- 1.4% |
| KO + WT | 0.113 | 0.053 | 748 | 0.001938 | -10.30 +/- 2.0% |
| KO + Juxta K | 0.047 | 0.048 | 547 | 0.002052 | -62.70 +/- 1.7% |
| KO + F349A | 0.11 | 0.069 | 1128 | 0.002054 | -12.70 +/- 2.0% |
| KO + Juxta K, F349A | 0.027 | 0.038 | 324 | 0.002111 | -78.60 +/- 1.7% |
